# The spliceosome component U1-70k loads nascent transcripts onto FVE/SGS3 via co-condensation to embark on RNA silencing

**DOI:** 10.1101/2025.04.22.650084

**Authors:** Taerin Oh, Kaiye Liu, Niankui Li, Changhao Li, Songxiao Zhong, Jiaying Zhu, Sascha Laubinger, Xiuren Zhang

## Abstract

Despite biochemical framework of RNA silencing has been studied well, how nascent transcripts enter RNA silencing remains poorly understood. Here we recovered U1-70k, a canonical spliceosome factor, as a novel component in posttranscriptional gene silencing (PTGS). We show that U1-70k interacts with SGS3 and FVE, two key components that supply RNA substrate for RNA-dependent RNA polymerase 6 (RDR6) to generate dsRNA. U1-70k promotes the loading of nascent RNA into the SGS3/FVE complex in the nucleus, which is subsequently translocated onto RDR6 in the cytosol. Importantly, U1-70k contains C-terminal low complexity (LC) domain and displays liquid-liquid phase separation (LLPS) pattern. U1-70k forms co-condensate with SGS3, another intrinsically disordered protein (IDP). The co-condensation is critical for PTGS as U1-70k LC variants display reduced SGS3 co-condensation, loss of RNA loading onto SGS3, and impaired RNAi function. Taken together, U1-70k channels nascent RNA to SGS3/FVE via its LLPS action to enter PTGS pathway.

## Introduction

PTGS plays important roles in the growth, development and environmental adaptation, such as viral defense in plants. PTGS can be guided via microRNA (miRNA) and small interfering RNA (siRNA) that are loaded into Argonaute (AGO) proteins to form RNA induced silencing complex (RISC) to cleave complementary transcripts or pause their translation.^1^ miRNA is generated from hairpin-structured primary precursors (pri-miRNAs) via a microprocessor complex that is minimally comprised of DCL1, double-stranded RNA binding protein (DRB1/HYL1) and a multifunctional protein Serrate (SE).^2–7^ Unlike miRNA, siRNA originates from single-stranded RNA (ssRNA) that is transcribed from endogenous and transgene loci, as well as viral RNA.^6^ One key step in the upper stream of siRNA pathway is the conversion of ssRNA into double-stranded (ds) RNA by RDR6 and suppressor of gene silencing 3 (SGS3). SGS3 forms homodimer, stabilizes transcripts, and recruits them into RDR6 for dsRNA production before the action of DCL2/4.^8–10^ We have recently reported that Flowering locus VE (FVE), a well-known epigenetic component, moonlights the RNA silencing pathway via promoting SGS3 dimerization, allowing SGS3 to bind to and stabilize more efficiently ssRNA/RDR6.^11^ At a sub-cellular level, siRNA synthesis takes place in siRNA body, a cytosolic membrane-less organelle that is believed to harbor SGS3, RDR6, and DCL4 among other components in siRNA pathway.^12^

Although different species of siRNAs share the same production machinery such as SGS3, RDR6 and DCL2/4, the initial steps of their biogenesis are different. One type of endogenous siRNAs, named trans-acting siRNAs (ta-siRNAs), originate from native non-coding transcripts that initially are cleaved by AGO1 and AGO7, guided by miR173 and miR390, respectively. In a similar manner, phased siRNAs are derived from protein-coding transcripts that are initially processed by AGO1 and a few species of miRNAs.^13^ In these scenarios, SGS3 bridges RISC to RDR6, and channels miRNA/RISC-cleaved transcripts to the RDR6-centered dsRNA production factory. By contrast, bulk of endogenous and transgene transcripts do not undergo prior miRNA/RISC action.^6,14^ However, how these transcripts are sorted in the nucleus and launched into the cytosolic RDR6 machinery for dsRNA production remained as a long-term mystery.

U1-70k is a core component of the spliceosome U1 small nuclear RNA ribonucleoprotein complex (snRNP) that participates in the first splicing event of nascent transcripts from RNA polymerase II (Pol II). Mammalian U1-70k comprises of N-terminal RNA-recognition motif (RRM) domain and C-terminal intrinsically disordered region (IDR) called low-complexity (LC) region.^15^ The RRM domain forms a complex with *U1 snRNA* and physically interacts with RNA Pol II subunits 2 and 12, enabling co-transcriptional splicing of nascent transcripts.^15,16^ The LC domain contains evenly distributed basic and acidic residues, resulting in a strong self-assembling tendency to undergo LLPS eventually being aggregated in mammals.^17^ Interestingly, U1-70k aggregation caused by LLPS property seems to be a driving force for Alzheimer’s disease (AD) in a splicing-independent manner.^18–21^ Whether U1-70k contributes to other types of posttranscriptional processing besides splicing is unknown and nor is whether the LLPS property of U1-70k engaged in other biological functions in addition to mammalian AD.

Here through an un-biased genetic screening, we identified *u1-70k* as a new mutant that has a compromised siRNA production from the transgene and endogenous loci. We found that U1-70k interacts with SGS3 and FVE in the nucleus and facilitates the ssRNA loading into SGS3/FVE, which in turn transport ssRNA from the nucleus to the cytoplasm. Interestingly, Arabidopsis U1-70k also displays LLPS, leading to self-assembling tendency of U1-70k.

Importantly, this property is indispensable for the U1-70k-SGS3 interaction, and its function in PTGS. Thus, this study reveals a non-canonical function of U1-70k in RNA silencing and provides new insight into how nascent transcripts are routed from the nucleus to the cytoplasm for dsRNA production in PTGS pathway.

## Results

### U1-70k has a non-canonical function in PTGS

We have recently reported a dual-screening system to systematically recover the genes that are involved in either miRNA or siRNA pathways.^11^ In this screening system, the parental line (E5-4) expresses a section of genomic *PHABULOSA* (*PHB*) (exon4, intron4, and exon5), which harbors the miR165/166 complementary sequence after splicing (Fig. S1A). The miR165/166 target site is flanked by *Green Fluorescent Protein* (*GFP*) and *Luciferase* (*LUC*) genes and driven by the native *AGO10* promoter (*P_AGO10_-GFP-PHB-LUC*).^11^ We performed EMS screening of E5-4 and obtained a dozen of *attenuated RNA silencing* (*ars*) mutants exemplified by *fve*.^11^

Here we characterized the second mutant, *ars2-1,* which showed significantly enhanced LUC signal and accumulated *LUC* transcripts compared with those in the parental line E5-4 (Fig. 1A and B). The *ars2-1* mutant was back-crossed with Col-0 to generate F2 mapping populations. We mapped the causative mutation to a 1.8-Mb interval on chromosome 3, which includes two candidate genes (Fig. S1B, C). We performed *ars2-1* complementation assays with the two candidate genes and found that *At3g50670 (U1-70k)* was able to restore the LUC signal to the parental level. These results indicated that the mutation in *U1-70k* was responsible for increased LUC activity in the mutant and this mutant was subsequently renamed *u1-70k-1* (Fig. 1C∼E, S1D).

**Fig. 1.**
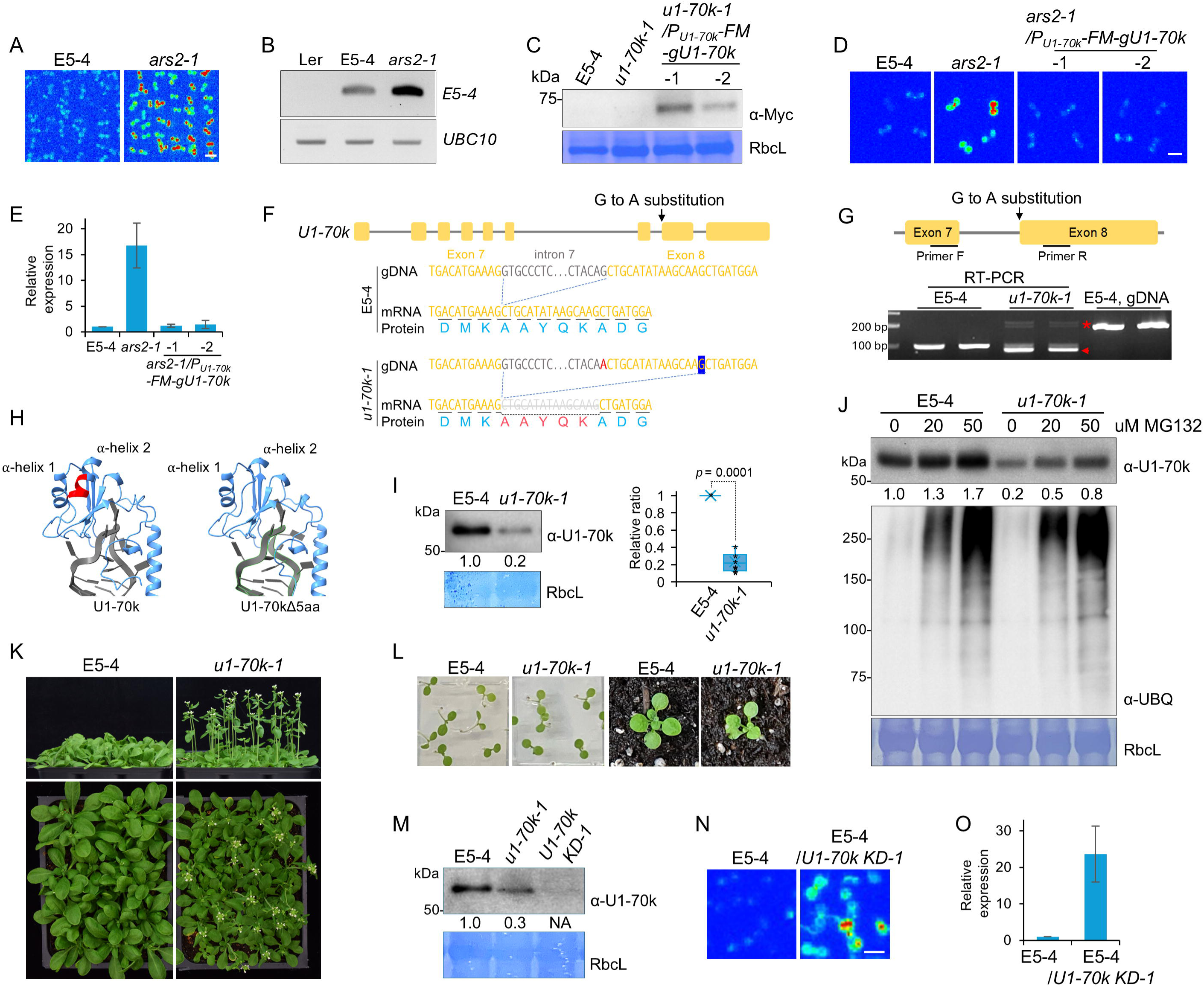
Genetic screening recovers U1-70k as a positive regulator in RNA silencing. (A) Enhanced LUC signal of 7-day-old seedlings of *ars2-1* vs E5-4. The image was captured under CCD camera for 240 seconds. Scale bars, 5 mm. (B) RT-PCR assay detected accumulated the reporter *GFP-PHB-LUC* transcript in *ars2-1* vs E5-4. *UBC10* was used for an internal control. (C) Western blot assays shows that *Flag-4Myc* (FM)-U1-70k accumulation in the complementation lines using an anti-Myc antibody. RbcL served as a loading control. (D) LUC assay shows that *pU1-70k-FM*-*U1-70k* could fully complement *ars2-1*. Scale bars, 5 mm. (E, O) Quantified LUC expression from Fig. 1D and I, respectively. Three replicates of LUC assay were quantified using image J. (F) Location of G to A substitution (red) in the junction of the 7^th^ intron and 8^th^ exon of *U1-70k* in the *u1-70k-1* allele and resultant changes in cryptic splicing, 15-nt skipping of mRNA and 5-aa deleted protein (red). Be noted: the cryptic G in blue box is used for splicing in *u1-70k-1.* (G) RT-PCR using a set of intron-flanked primers identified the 15-nt deleted (red arrowhead) and intron-retained isoforms (asterisk) of *U1-70k* transcript in *u1-70k-1* vs E5-4. (H) AlphaFold3 modeling predicts that the five-aa (red colored) deletion in *u1-70k-1* disrupts IZ-helix 2 in the RRM domain. (I) Western blot assay, followed by statistical analysis, demonstrates reduced accumulation of endogenous U1-70k variant in *u1-70k-1* vs WT protein in E5-4 controls. The relative ratio of U1-70k protein was quantified using image J, based on data from seven independent western blot assays. (J) Western blot assay shows that U1-70k and U1-70kΔ5aa are subjected to the 26S proteasome regulation. U1-70k and poly-ubiquitin chains of total proteins was detected by antibodies against endogenous U1-70k and ubiquitin respectively. RbcL serves as a loading control and the relative band intensity was measured by image J (I, J). (K, L) Growth and developmental phenotypes of *u1-70k-1* and E5-4 in the stages 3.5-week-old plants (K), and 7-day and 2-week-old plants (L). (M) The endogenous U1-70k was barely detectable in the *U1-70k* KD line. RbcL serves as a loading control and the relative band intensity was measured by image J. (N) CCD imaging showed increased LUC signal in *U1-70k* KD plants vs WT plants. Scale bars, 5 mm.

The *u1-70k-1* mutation is a G to A substitution at the end of intron 7 in the *U1-70k* gene, which activated a cryptical 3’ splicing site, leading to 15-nucleotide (nt) deletion in 5’ region of exon 8, and resultant five-amino acid (aa) deletion in the U1-70k protein (Fig. 1F). The 15-nt-deleted *U1-70k* transcript variant was predominantly expressed in *u1-70k-1* but minor portions of wild type and intron-retained *U1-70k* transcript were also observed in the mutant (Fig. 1G).

The five-aa deleted region was located on α-helix 2 (αH-2) in the RRM domain, which otherwise stabilizes a β-sheet motif that physically interacts with U1 snRNA (Fig. 1H, S1E). This region has a highly conserved sequence across diverse species of mammals and plants, suggesting that the absence of the five aa may lead to a defect in U1-70k function (Fig. S1F). Western blot results using a homemade U1-70k antibody demonstrated that the 5-aa deleted U1-70k variant (refer as U1-70kΔ5aa) was less stable than wild-type U1-70k, due to its quick degradation by 26S-proteasome (Fig. 1I, J, S1G, H). This result suggested that the *u1-70k-1* mutation should be a recessive loss-of-function mutation. However, *u1-70k-1* was morphologically undistinguished from the WT plants (or only showed subtle growth abnormality) in the seedling stage (Fig. 1L). Only at the adult stage, *u1-70k-1* lines displayed a short stature and an early flowering phenotype but with normal silique development (Fig. 1K).

We further created the *U1-70k* knockdown (KD) plants via an artificial miRNA strategy (Fig. 1M, S1H, I).^4^ Again, *U1-70k* KD lines showed increased LUC signal similar to that of *u1-70k-1* (Fig. 1N, O). This result further validated that U1-70k is a *bona fide* component in RNA silencing pathway. Intriguingly, *U1-70k* KD lines showed severe defective growth exemplified by bushy growth and clustered phyllotaxy of siliques, reminiscent of the phenotype observed in a previous report (Fig. S1J).^22^ These results indicated that *U1-70k* KD lines were much more severe mutants than the *u1-70k-1* allele, which was just a hypomorphic allele. Expected from the phenotypic difference, *U1-70k* KD and *u1-70k-1* might regulate splicing and silencing processes differently (Fig. 1K, L, S1J).

### U1-70k functions in upstream of siRNA pathway

Next, we attempted to pinpoint the action site of *U1-70k* in RNA silencing via sRNA sequencing (Fig. S2A). To update, 147 of earlier proposed 326 miRNA species have been re-annotated as *bona fide* ones (DCL1-dependent).^23^ Among these, the predominant number of miRNAs did not show obvious changes in their expression levels (Fig. 2A). This result was validated via sRNA blot assays with a few found miRNAs (Fig. S2B). In lines with this, qRT-PCR assays did not detect significant change of the expression of miRNA targets between *u1-70k-1* and WT plants (Fig. S2C). Since the E5-4 reporter is driven by *AGO10* promoter, we measured the expression of *AGO10* (Fig. S2C). We observed that *AGO10* expression was marginally upregulated, but this increase was not comparable to the one of *LUC* mRNA in *u1-70k-1* vs WT (Fig. S2C). This result indicated that highly accumulated *LUC* transcript in *u1-70k-1* might not be through the transcriptional gene silencing (TGS) pathway. Taken together, we tended to interpret that the *u1-70k-1* mutation does not seem to affect miRNA or TGS pathways, with a further suggestion that the mutation might impact the siRNA-mediated PTGS pathway.

**Fig. 2.**
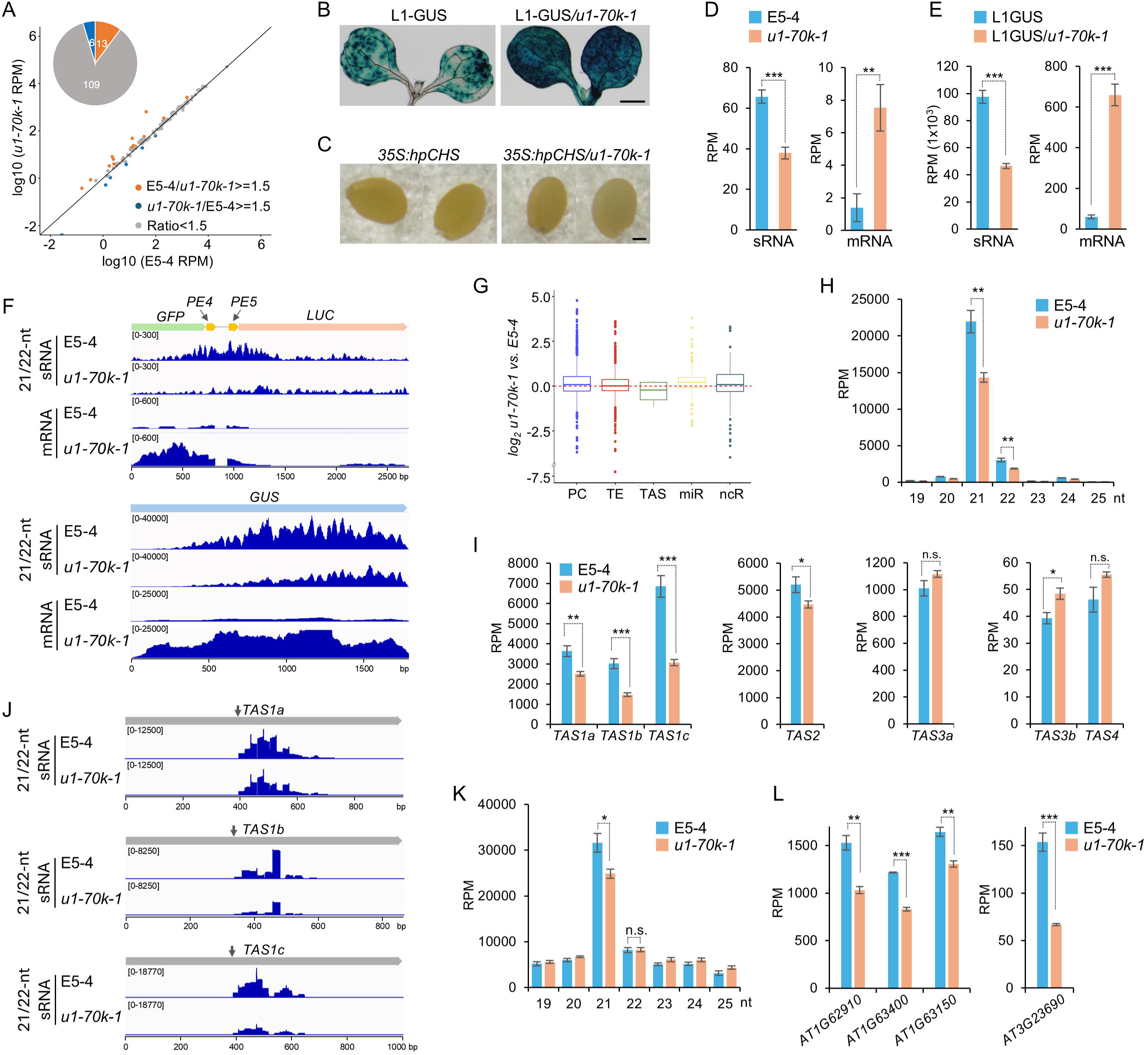
U1-70k promotes biosynthesis of siRNAs from sense transgene reporters and endogenous loci. (A) sRNA-seq analysis showed the majority of canonical miRNAs were not altered in *u1-70k-1 vs* E5-4. The x and y axis indicate the logarithms of normalized expression levels of miRNAs in WT and *u1-70k-1*, respectively. (B) GUS histochemical staining shows that *u1-70k-1* causes a silencing defect in L1 reporter line expressing sense GUS gene. The 7-day-old seedlings of L1 GUS and L1 GUS/*u1-70k-1* lines were used for assays. Scale bar, 1 mm. (C) The analysis of dsRNA-mediated silencing of *CHS* reporter line, indicated by the white seed color, does not suggest that *u1-70k-1* impacts downstream of PTGS pathway. Scale bar, 0.1 mm. (D, E) mRNA-seq and sRNA-seq show that *u1-70k-1* reduced siRNA whereas increased mRNA amount of the two independent reporter genes *LUC* (D) and *GUS* (E) in the mutant vs E5-4 and L1 controls, respectively. (F) The Integrative Genomics Viewer (IGV) graphs show the reads distribution of 21-to 22-nt sRNAs and mRNA in the *GFP-PHB-LUC* and *GUS* gene bodies. (G) A box plot analysis shows that the accumulation of ta-siRNAs but no other species of sRNAs is clearly reduced in *u1-70k-1* vs E5-4. Protein coding (PC), transposon (TE), TAS, miRNA (miR) and non-coding RNA (ncR). (H, K) Bar plots show that the pattern changes of ta-siRNA (H), phased siRNA (K) in *u1-70k-1* vs E5-4. (I) Computational analysis shows that AGO1-miRNA-dependent ta-siRNA from *TAS1a∼c* and *TAS2*, but not AGO7-miR390 dependent ta-siRNA from ta-siRNA *TAS3a/b* (I), is reduced in *u1-70k-1* vs WT. (J) IGV graphs shows the reads distribution of 21- to 22-nt sRNAs in the *TAS1* RNAs’ gene bodies. The arrow indicates the AGO1/miR173 cleavage site. (L) The phased siRNA from PPR genes processed by AGO1-miR161 (left graph) and *AT3G23690* cleaved by AGO1-miR393 (right graph) is reduced in *u1-70k-1* vs WT. The asterisks indicate *p*-value from student’s *t*-test (***, *p* < 0.001; **, *p* < 0.005; *, *p* < 0.02).

To further test this, we crossed *u1-70k-1* with the L1 line, which contains a silenced *ß-glucuronidase* (*GUS*) gene and has been broadly used as the reporter system for the sense-transgene PTGS pathway.^24^ The homozygotes of L1-GUS*/u1-70k-1* showed obviously enhanced GUS signal compared with the parent control, indicating that *U1-70k* indeed regulates sense transgene silencing (Fig. 2B). In parallel, we crossed *u1-70k-1* with the *35S:hpCHS* reporter line that the expresses *hpCHS* (*hairpin chalcone synthase*) and typically serves as a reporter for the downstream pathway of PTGS.^25^ *CHS* synthesizes flavonoid pigments, resulting in the dark-brown seed color in WT plants, whereas dsRNA of *CHS* causes its silence, causing light seed color. Here, the *u1-70k-1* mutation did not recover the white color *of 35S:hpCHS* seed to the brown one in WT seeds (Fig. 2C). The result indicated that U1-70k may be involved in dsRNA synthesis or earlier steps, but not in the downstream in sense transgene PTGS pathway (Fig. 2B, C).

We performed RNA-seq for *u1-70k-1* vs WT (Fig. S2D, E). In lines with the dramatic increase in LUC and GUS signals in the mutant, RNA-seq also showed the steady-state levels of *LUC* and *GUS* transcripts were increased by 7 and 10 times in *u1-70k-1* vs WT controls, E5-4 and L1, respectively (Fig. 2D∼F). By contrast, sRNA-seq revealed that the reads of 21/22-nt sRNA mapped with *E5-4* and *L1 GUS* loci were decreased more than 40% in the mutant vs WT plants (Fig. 2D∼F). Together, we concluded that U1-70k has a positive regulatory function in the upstream of sRNA biogenesis, prior to the dsRNA cleavage by DCL2/4.

### U1-70k promotes production of ta-siRNAs and phased siRNAs

We next investigated if U1-70k impacted sRNA production from endogenous loci on a genome-wide scale. Endogenous sRNAs can derive from the transcripts of protein-coding genes, transposon elements (TE) and non-coding regions.^14^ The box-plot analysis of sRNA-seq did not reveal the substantial changes of the endogenous siRNAs (Fig. 2G). *TAS* loci transcribe species of non-coding transcripts that can be initially cleaved from RISC activity and subsequently serve as templates for RDR6 to produce dsRNA and eventually ta-siRNAs.

Interestingly, the total reads number of 21/22-nt sRNAs mapped on the *TAS* loci were clearly decreased in *u1-70k-1* vs *E5-4* (Fig. 2G). Further computational analysis detected approximately 36% and 41% decreases for 21 and 22-nt sRNA from the *TAS* loci, respectively, in the mutant vs WT plants (Fig. 2H). These results indicated that U1-70k impacted ta-siRNA production.

Ta-siRNAs can be further classified into three categories: ta-siRNA from *TAS1a-c* and *TAS2* result from the “one-hit” activity of miR173/AGO1-centered RISC on *TAS1* precursors. Similarly, *TAS4* is derived from the substrate cleaved once by miR828/AGO1-RISC.^26^ By contrast, ta-siRNA from *TAS3a/b* is processed from the “two-hit” action of miR390/AGO7 on *TAS3* substrates.^27^ Interestingly, the ta-siRNAs from *TAS1a-c* and *TAS2* were significantly reduced in *u1-70k-1* vs WT, whereas *TAS3a* and *3b* ta-siRNAs were not (Fig. 2I, J). These results suggested that U1-70K might positively regulate the miR173/AGO1-initiated pathway. Despite dependence on AGO1 activity, ta-siRNA from *TAS4* was not decreased, likely due to low counts of the siRNAs.

Extended from ta-siRNAs are phased siRNAs that also entail the initial cleavage of AGO1-miR161 and miR393 of protein-coding transcripts prior to dsRNA production.^13,28^ Again, the amount of phased siRNAs was also reduced significantly in *u1-70k-1* vs WT (Fig. 2K).

Especially, the expression of phased siRNAs, whose synthesis begins from miRNA/AGO1 cleavage, is clearly reduced. (Fig. 2L, S2F). In all, the production from ta-siRNA and phased siRNAs generally minored the production of sense PTGS siRNA in *u1-70k-1* vs WT. We interpreted that U1-70k could affect siRNA production, at least from ta-siRNA and phased siRNA loci.

### U1-70k promotes siRNA biogenesis independently from its canonical role in splicing

Since U1-70k has a canonical function in U1 snRNP in splicing, we wondered if the role of U1-70k in gene silencing is related to U1 snRNP or splicing. To test this, we crossed mutants of *u1a*, which encodes a partner of U1x-70K in U1 snRNP complex, with Ec5-4 (referring E5-4, back crossed with Col-0 for seven times).^15^ In parallel, we also obtained a homozygous allele of Ec5-4 crossed with the triple mutant of *luc7a/b/rl* which encode three other auxiliary components of U1 snRNP and showed severe pleotropic defects in development.^29^ In addition, we created a KD mutant in the E5-4 background for *U1C*, encoding another component in U1 snRNP (Fig. S2G. H).^15^ These factors are components of the U1 snRNP and are closely related to the U1-70k function in splicing. We found that *luc7 a/b/rl* triple mutants displayed slightly enhanced, but less than 2-fold, LUC signal (Fig. 3A). However, *u1a* mutants or *U1C-amiR* KD plants did not show stronger LUC compared with their cognizant controls (Fig. 3B, C). These results indicated that the function of U1-70k in PTGS could be separated from its canonical activity in U1 snRNP complex.

**Fig. 3.**
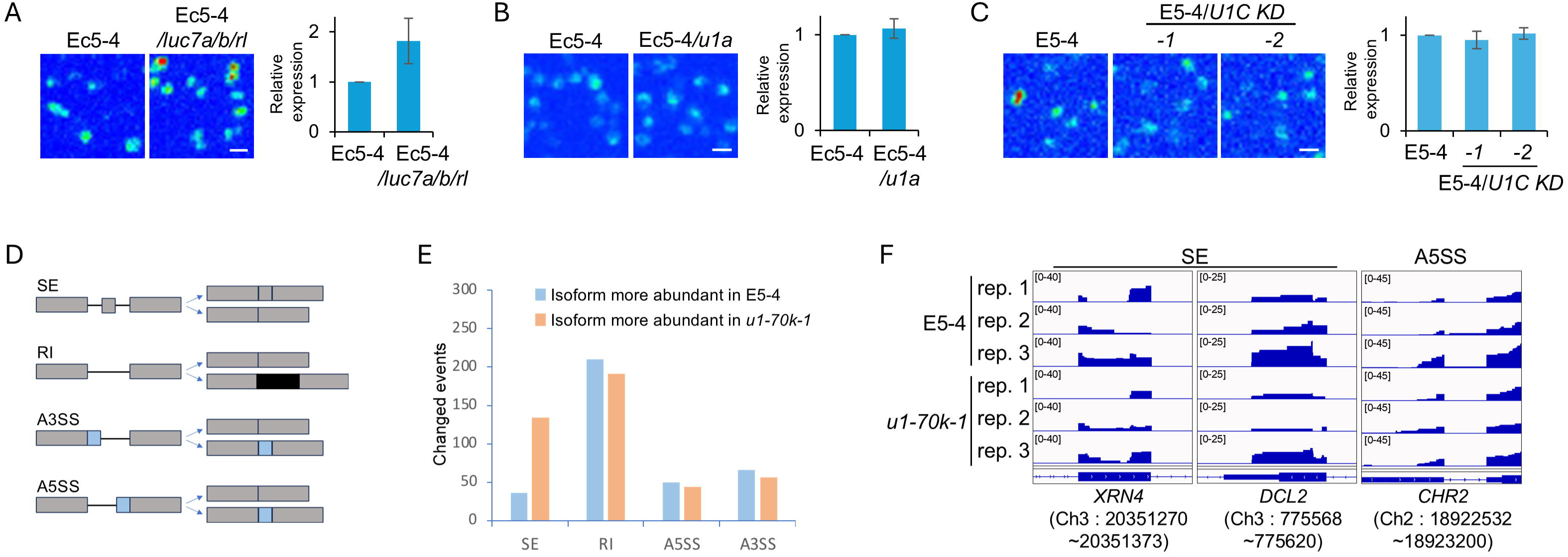
U1-70k promotes siRNA biogenesis independently from its canonical role in splicing. (A∼C) LUC imaging shows that loss-of-function of the genes encoding U1-70k accessory proteins *LUC7a/b/rl, U1A, U1C,* could not release PTGS when introduced into the E5-4 reporter line. Quantification of LUC signal from three replicates via image J is shown on the right. Scale bars, 5 mm. (D) Schematic diagrams demonstrate four different types of alternative splicing such as skipped exon (SE), retained intron (RI), alternative 3’ splice site (A3SS) and, alternative 5’ splice site (A5SS). (E) RNA-seq analysis shows that the numbers of alternative splicing events in E5-4 and *u1-70k-1.* (F) The IGV graphs did not reveal significant differences in the reads distribution of mRNA on the alternative splicing site in *XRN4, DCL2* and *CHR2*.

We then assessed the impact of *u1-70k-1* on splicing events. RNA-seq could detect some difference of splicing defects between *u1-70k-1* and WT plants, exemplified by increased skipping exon (SE) events but marginally decreased retained-intron (RI) events in the mutant vs WT (Fig. 3D, E). However, these aberrant splicing events might be considered as a minor effect at a global level when compared to the *luc7a/b/rl* mutant, which displayed substantially increased of SE and IR transcripts compared to WT.^29^ The different impact of U1-70k and LUC7a/b/rl on splicing was likely because *u1-70k-1* is a hypomorphic allele whereas the *luc7a/b/rl* mutant is a null allele. Given that mutations in *U1A* or *U1C*, or even *LUC7a/b/rl* has defects in U1 snRNP function in splicing but did not release *LUC* silencing in the E5-4 report line, we tended to believe that the defect in RNA silencing might not be through an indirect splicing defect in *u1-70k-1* vs WT.

Subsequently, we focused on the expression or splicing patterns of transcripts of core components or auxiliary regulators in various sRNA and RNA decay pathways. Among 80 known factors, none of them showed significantly different expression in *u1-70k-1* vs WT (Fig. S2I). Computational analysis suggested the presence of skipped-exon events, or usage of an alternative 5’ splice site for *XRN4*, *DCL2*, and *CHR2*, which participate in RNA decay, dsRNA processing, and miRNA biogenesis, respectively (Fig. 3E). However, Integrative Genomics Viewer (IGV) did not show obviously contrasted splicing patterns of the transcripts in *u1-70k-1* vs E5-4 (Fig. 3F). Overall, these results suggest that the reduced RNA silencing in *u1-70k-1* did not result from downregulation or misprocessing of transcripts that code for RNA silencing components. In all, we concluded that U1-70k can promote siRNA biogenesis independently from its canonical role in pre-mRNA splicing.

### U1-70k interacts with SGS3/FVE in nucleus

We next hypothesized that U1-70k might function via partnering with some component(s) in PTGS pathway. To test this, we performed yeast two-hybrid screening (Y2H) for a handful of candidates in the pathway. Such Y2H screening recovered SGS3 and FVE as the partners of U1-70k (Fig. 4A, S3A). Earlier, we reported that FVE can be localized in both the nucleus and cytoplasm and its function in the cytoplasm can be uncoupled from its role in the epigenetic silencing in the nucleus.^11^ Here, in lines with this result, the colocalization of U1-70k and FVE was readily detected in the nuclei in confocal microscopy imaging when the genes were co-expressed in tobacco leaves (Fig. 4B). Furthermore, the U1-70k/FVE interaction was also validated with bimolecular fluorescent complementation (BiFC) assays when the c-YFP-tagged U1-70k was co-expressed with cYFP-tagged FVE in *N. Benthamiana* (Fig. 4C, S3B).

**Fig. 4.**
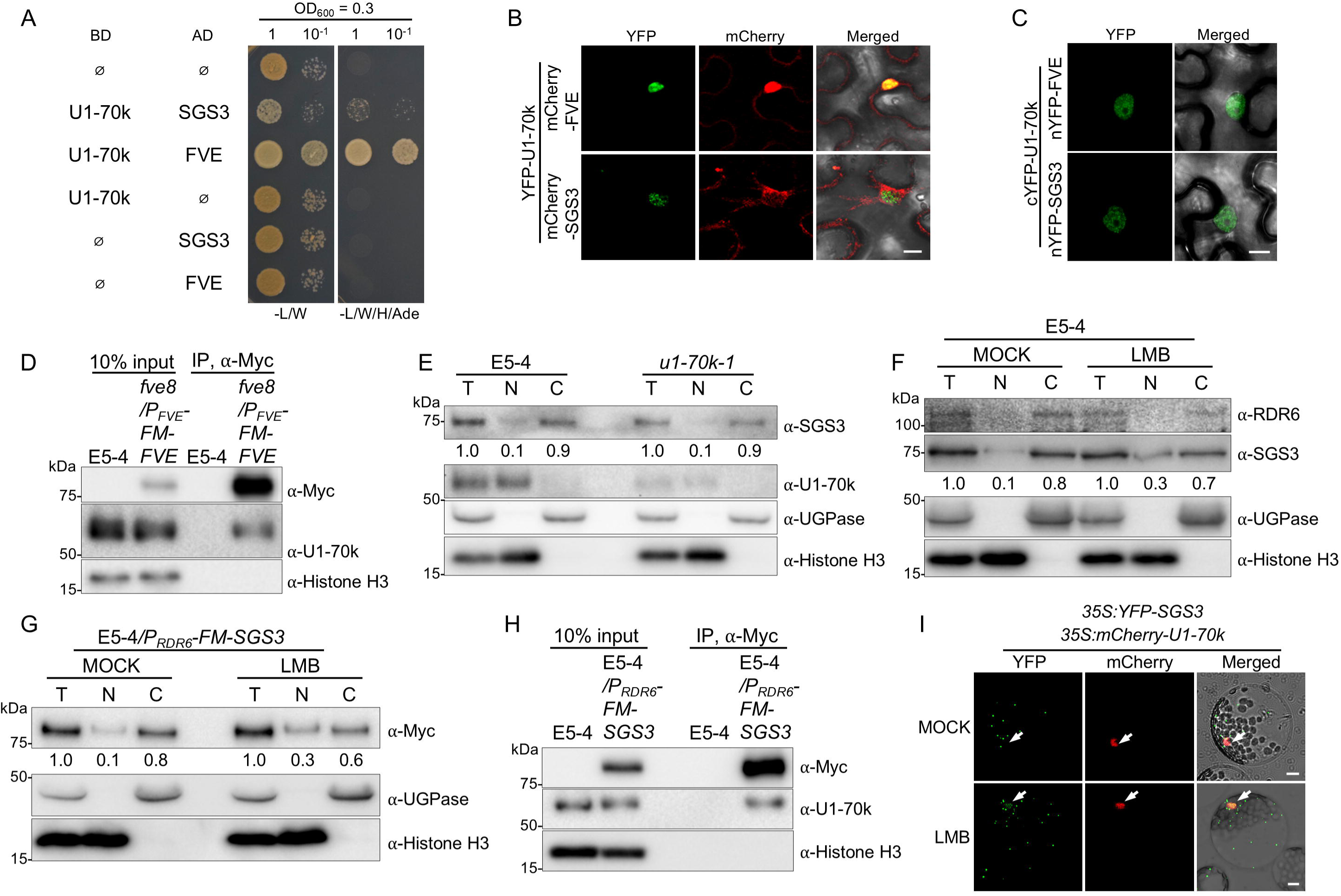
U1-70k interacts with SGS3 and FVE in the nucleus. (A) Y2H screening recovered SGS3 and FVE as partners of U1-70k. AD, activation domain of galactose-responsive transcription factor (GAL4); BD, DNA binding domain. (B) Confocal microscope images showed that U1-70k obviously colocalized with FVE but marginally with SGS3 in the nucleus of tobacco cells. (C) BiFC assay indicated that cYFP-tagged U1-70k could complement nYFP-SGS3 and -FVE in tobacco cells. Scale bars in B and C indicate 5 µm. (D) Western blot assay showed that U1-70k was detected in FVE immunoprecipitants. (E) Western blot assays of nuclear/cytosol fractionation showed that a small portion of SGS3 occurred in the nuclear fraction. (F, G) Western blot assays showed that SGS3 was more accumulated in the nucleus upon leptomycin B (LMB) treatment. UGPase and H3 served as cytosolic (C) and nuclear fraction (N) marker, respectively. T, total crude extracts. The relative band intensity was measured by image J (E∼G). (H) Western blot assays showed that U1-70k was co-purified with SGS3 from the Co-IP assay using the LMB-treated nuclear fraction. In Panel D and H, IP was performed using an anti-Myc, Co-IP products were detected using an anti-U1-70k. H3 serves as a negative control. (I) Confocal image showed that YFP-SGS3 and mCherry-U1-70k were co-localized in the nucleus in Arabidopsis protoplast under LMB treated condition. Scale bars, 10 µm.

Importantly, we immunoprecipitated Flag-4Myc-FVE (FM-FVE) from a stable complementation line expressing *fve-8; P_FVE_-FM-FVE* and could clearly detect U1-70k in the co-immunoprecipitant (Fig. 4D). This interaction should take place in the nucleus as U1-70k is not localized in the cytoplasm whereas FVE can shuttle between the nucleus and cytoplasm.

We could not detect the co-localization of U1-70k and SGS3 in the confocal microscope imaging, which may be due to SGS3 predominantly localizing in the cytosol, while U1-70k is present in the nucleus (Fig. 4B). However, we observed the YFP signal in the nucleus from nYFP-SGS3 and cYFP-U1-70k in the BiFC assays, indicative of their interaction in the nucleus (Fig. 4C, S3B, C). Then, we hypothesized that SGS3 might shuttle between the cytosol and the nucleus and interact with U1-70k in the nucleus. To test this idea, we conducted a nuclear-cytosol fractionation assay. While SGS3 was mainly detected in the cytosolic fraction, we could indeed observe a small portion of SGS3 from the nuclear fraction (Fig. 4E). This result suggested that SGS3 might be a nuclear/cytosol shuttling protein.

To further test this, we applied to E5-4 plants with a nuclear export inhibitor, leptomycin B (LMB), which blocks Exportin 1/Chromosomal Region Maintenance 1 (EXP1/CRM1) mediated nuclear export.^30^ Indeed, upon the LMB treatment, SGS3 could be more easily detected in the nucleus (Fig. 4F, G). Then we carried out the Co-IP assay with the LMB-treated nuclear fraction. Notably, U1-70k was co-purified with SGS3 in this scenario (Fig. 4H). In lines with this, we could detect nuclear SGS3 co-localized with U1-70k in Arabidopsis protoplast under LMB treated condition (Fig. 4I, S3D). Overall, we proposed that SGS3 is a nuclear/cytosol shuttling protein and can interact with U1-70k in the nucleus.

RDR6 has been detected in the nucleus in trichome cell and root tips of Arabidopsis seedling or when RDR6 was overexpressed in tobacco leaves, here RDR6 was mainly detected in the cytosol fraction in two-week-old seedlings.^31–33^ This result is consistent with other previous studies that show cytosol and sRNA body localization of RDR6 in tobacco leaves and Arabidopsis mesophyll cells (Fig. 4F, S3E, F).^12,34–37^ One possibility for our failure to detect the nuclear function of RDR6 might be due to the relatively low sensitivity of the anti-RDR6 antibody in our hands (Fig. 4F, S3E, F).

Interestingly, *u1-70k-1* codes the five-aa deleted U1-70k variant in mutant. Y2H assays showed that the U1-70kΔ5aa still interacted with SGS3 and FVE as well as WT U1-70k (Fig. S3G). This result suggested that the compromised function of U1-70kΔ5aa in PTGS should result from a mechanism different from its partnership with SGS3/FVE in the mutant vs WT.

### U1-70k promotes ssRNA loading into SGS3 in vivo

We further pinpointed the cellular compartments where the PTGS was mainly compromised in *u1-70k-1*. To this end, we conducted RT reaction with an oligomer dT primer followed by PCR assays separately with nuclear/cytosol fractionations. We detected marginally matured *GFP-PHB-LUC* transcript (indicated by an arrowhead) in the nuclear fraction from E5-4 but approximately two-fold accumulation from *u1-70k-1* vs WT (Fig. 5A, B). In the cytosolic fraction the reporter transcript was silenced and beyond detectable limit by RT-PCR for E5-4, while the transcript was obviously detected for *u1-70k-1* (Fig. 5B). The intron-containing reporter transcript (indicated by an asterisk) were more accumulated in the total and nuclear fractions, each with about two- and five-fold increase, respectively, but not in the cytosolic fraction, from *u1-70k-1* vs WT (Fig. 5B). These results suggested that the nascent (or not fully processed) transcript of the E5-4 reporter remains to be in the nucleus in the mutant whereas the portion of transcript is trafficked out of the nucleus and then consumed in the cytosol in WT plants (Fig. 5B).

**Fig. 5.**
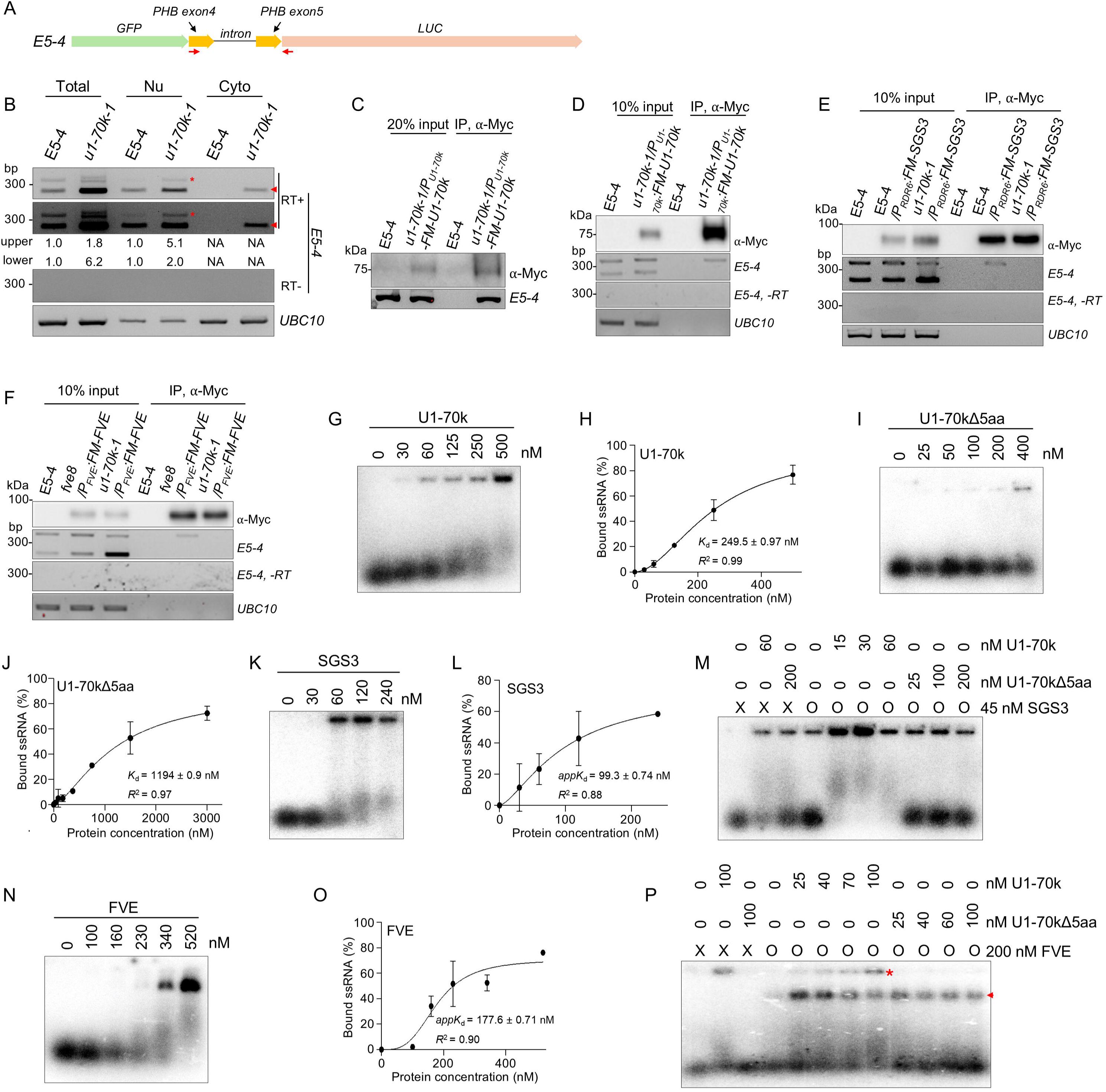
U1-70k promotes nascent RNA loading into SGS3/FVE in vivo and in vitro. (A) Binding location (red arrowheads) of the primer set used for RT-PCR in the *E5-4* reporter transcript in Fig. 5B∼E. (B) RT-PCR showed that silencing of *GFP-PHB-LUC* transcript predominantly occurred in the cytosol (Cyto) and was inhibited in *u1-70k-1*. An oligo dT primer was used for RT reaction. The asterisk and arrowhead indicate nascent *GFP-PHB-LUC* transcript and matured transcript, respectively. The relative band intensity was measured by image J. (C) ChIP-PCR assay demonstrated that U1-70k clearly bound to the *GFP-PHB-LUC* locus in the complementation lines. (D) RIP-PCR result showed that U1-70k could bind with nascent *E5-4* transcripts. (E, F) RIP-PCR assays exhibited that nascent, but not matured, *GFP-PHB-LUC* transcript was readily detectable in SGS3 (E) and FVE (F) immunoprecipitants in E5-4, but not in *u1-70k-1*. ChIP or RIP was performed using the nuclear fractions of the LMB treated seedlings using an anti-Myc antibody and *UBC10* served as an internal control (C∼F). (G∼J) EMSA assays demonstrated that U1-70k binds to ssRNA at a higher affinity than U1-70kΔ5aa, which carries 5-aa deletion in the RRM domain. (K, L, N, O) EMSA results showed that SGS3 (K) and FVE (N) could bind to 38-nt long homogenous ssRNA. (H, J, L, O) The binding affinities (*K*_d_) of protein/ssRNA were calculated from quantification of Fig. 5G, I, K, N and S5B, D. (M, P) The EMSA assay demonstrated that U1-70k could clearly enhance ssRNA loading onto SGS3 (M) and FVE (P), whereas U1-70kΔ5aa does not. In the panel (5P), the asterisk and arrowhead indicate U1-70k/ssRNA and FVE/ssRNA complexes, respectively.

We repeated RT-PCR assay above but with random primers for RT (Fig. S4C, left set). RT-PCR with a random primer amplifies either nascent RNA prior to polyadenylation or intron-retained mRNA but with poly A tail. In this scenario, we detected comparable amount of a total of nascent and intron-retained *LUC* reporter transcript in the nuclear fraction of E5-4 and *u1-70k-1* (Fig. S4C, left set). The contrasted RT-PCR results using two different sets of RT primers indicated that a large portion of nascent transcripts are routed out to RNA silencing pathway before polyadenylation of pre-mRNA in WT plants, while the nascent portion of transcripts does not escape from the nucleus and then is degraded in *u1-70k-1* vs WT.

Given that that mammalian U1-70k physically interacts with RNA Pol II and that U1-70k can also interact with SGS3 and FVE in the nucleus, we speculated that nuclear U1-70k/SGS3/FVE may recruit nascent transcripts co-transcriptionally on the *E5-4* reporter locus.^16^ Indeed, ChIP-PCR could detect the accumulation of U1-70k on the E5-4 locus (Fig. 5C, S4D). However, SGS3 was not enriched in the locus (Fig. S4D, E). These results suggested that U1-70k but not SGS3 is associated with the transgene locus, with a further suggestion that U1-70k may recruit the nascent RNA from the E5-4 locus co-transcriptionally in chromatin and then transfer the transcript to SGS3 in nucleoplasm. To test this, we performed RIP-PCR assays in the E5-4 and *u1-70k-1* background. Interestingly, the nascent *E5-4* transcript could be readily detected in the U1-70k immunoprecipitants but not in the control IP from E5-4 (Fig. 5D).

Furthermore, the nascent RNA could be easily recovered from either SGS3 or FVE RIP (Fig. 5E, F, S4F). By contrast, we did not detect the nascent E5-4 transcripts in SGS3 and FVE RIP in the *u1-70k-1* background (Fig. 5E, F, S4F). These results indicated that U1-70k indeed absorbed the nascent RNA and subsequently loaded them into SGS3/FVE in the nucleus.

Importantly, the same results were obtained for the L1 reporter line (Fig. S4G). Since *GUS* transcript in the L1 line, different from *GFP-PHB-LUC* RNA, does not contain an intron, U1-70k recruitment of the nascent transcripts does not depend on its association with U1 snRNP. Taken together, we concluded that U1-70k can bind to nascent transcripts and channel their loading into SGS3 and FVE in the nucleus.

### U1-70k enhances SGS3/FVE binding of transcript in vitro

We next investigated biochemical feature of U1-70k and its possible impact on the SGS3/FVE-RNA interaction. To this end, we conducted electrophoretic mobility shift assay (EMSA) in vitro (Fig. S5A). U1-70k binds to the ssRNA with *K*_d_ = ∼ 250 nM, while U1-70kΔ5aa showed dramatically reduced affinity to ssRNA (*K*_d_ = ∼ 1200 nM, Fig. 5G∼J, S5B∼D). These results indicate that the five-aa deletion right in the RRM domain significantly reduced the association of U1-70k with RNA substrates, resulting in compromised function of U1-70k, which in turn triggered its quick degradation by 26S proteasome (Fig. 1H, L, 5I, J, S5B∼D).

SGS3 binds ssRNA with *K*_d_ = 100 nM. This value is higher than the ones in previous findings (Fig. 5K, L, S5D), and this discrepancy may be attributed to the use of a different RNA substrate in the current experiment.^11^ Interestingly, the co-incubation of 45 nM SGS3 and 15 nM U1-70k showed a synergistic property of RNA binding (Fig. 5M, S5E). In contrast, application of 200 nM U1-70kΔ5aa, which was 13 times the concentration of U1-70k, did not show such effect (Fig. 5M, S5E). Thus, both in vitro EMSA and in vivo RIP results indicated that U1-70k could strongly enhance the RNA binding activity of SGS3. We repeated EMSA assays with FVE with U1-70k or U1-70kΔ5aa variant. Again, we found that U1-70k, but not U1-70kΔ5aa, could also elevate the RNA binding of FVE, reminiscent of cooperation of U1-70k with SGS3 (Fig. 5N∼P, S5D). Taken together, we concluded that U1-70k could directly interact with SGS3/FVE and promote their RNA binding. These results were well aligned with prerequisite of U1-70k for SGG3/FVE recruitment of nascent transcripts in vivo (Fig. 5E, F, S4F, G).

### Self-assembly of U1-70k is critical for the SGS3 interaction

U1-70k is an IDP with an RRM in the middle flanked with N- and C-terminal IDR regions, each of approximately 50 aa and 200 aa, respectively (Fig. S6A). The N-terminal IDR is unique to plants, suggesting that it may serve a plant-specific function (Fig. S6B). In the C-terminal IDR region, the LC1 region is enriched with basic/acidic amino acid-repeated sequences in both mammals and plants, while the LC2 domain exhibits sequence variation, highlighting the potential importance of LC1 domain (Fig. S6C). Further Y2H assays showed that FVE interacted with N-terminal region of U1-70k, whereas SGS3 did not interact with either N- or C-terminal part of U1-70k (Fig. 6A∼C). This result suggested that the SGS3 interaction with U1-70k might entail its conformation integrity.

**Fig. 6.**
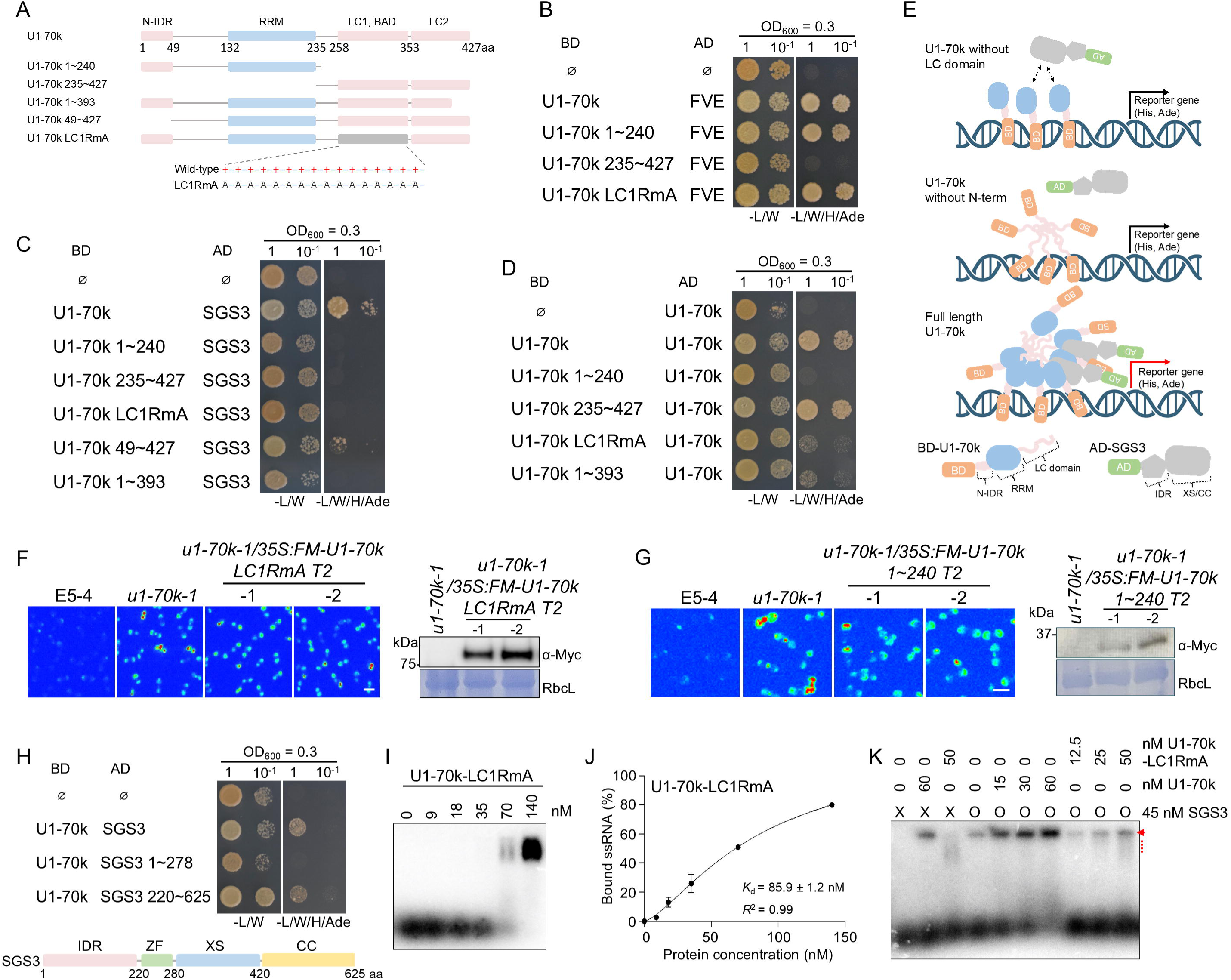
The IDR domain of U1-70k is critical for its interaction with SGS3 and RNA loading. (A) Schematic constructs of U1-70k variants used in the Y2H assays. The gray color indicates LC1 domain with R to A substitution (LC1RmA). (B) Y2H assay shows that the N-terminal region of U1-70k interacts with FVE. (C) Y2H assay shows that the full-length, N-terminal region, and LC domains of U1-70k are necessary for SGS3 interaction. (D) Y2H assay indicates that the LC domain drives self-assembly of U1-70k. (E) A proposed model demonstrates that the full-length conformation and self-assembly properties of U1-70k are essential for SGS3 interaction in yeast. Top: SGS3 cannot activate reporter genes due to its weak binding affinity for the U1-70k N-term region. Middle: The assembled LC domain does not interact with SGS3. Bottom: The full-length assembled U1-70k binds to SGS3, results in activation of the reporter genes. (F, G) Complementation assays show that deletion or R to A substitutions in the LC domain disrupted U1-70k’s function in PTGS. Scale bars, 5 mm. Western blot assay of the complementation lines using an anti-Myc antibody and RbcL served as a loading control. (H) Y2H assay demonstrated that U1-70k interacts with the truncated SGS3 without N-terminal IDR and zinc finger domain (ZF). Bottom: A schematic structure of different domains of SGS3. (I) EMSA assay demonstrates that U1-70k-LC1RmA had enhanced ssRNA binding affinity vs U1-70k. (J) The binding affinity (*K*_d_) of U1-70k-LC1RmA/ssRNA was calculated from quantified EMSA result from panel J. (K) EMSA shows that U1-70k-LC1RmA did not promote ssRNA loading into SGS3. The arrowhead and dashed line indicate U1/70k/SGS3/ssRNA and U1-70k-LC1RmA/ssRNA complexes, respectively.

We noticed that the lengthy LC region of mammalian U1-70k tends to promote self-assembly and also undergoes LLPS.^17^ Then, we conceived an idea that Arabidopsis U1-70k might also have similar features that may be crucial for its interaction with SGS3. Indeed, Arabidopsis U1-70k clearly showed self-interaction (Fig. 6D, S3H). This self-interaction was through its C-terminal LC domain as U1-70k (235-427aa), which harbors the LC domain alone, could maintain its self-association, whereas the absence of the LC domain destabilized U1-70k self-interaction (Fig. 6D, S3H). Notably, the partially deleted LC2 (394∼427aa) U1-70k and U1-70k-LC1RmA, in which all arginine were mutated to alanine in the LC1 domain, displayed dramatically weakened self-interaction in Y2H assays, indicating that the integrity of LC domain is essential for its self-assembly (Fig. 6D, S3H). Next, we investigated how the self-assembly of U1-70k impacted its interaction with SGS3. To this end, we performed Y2H assays with a series of new U1-70k variants.

Interestingly, the N-IDR-deleted U1-70k (U1-70k (49∼427aa)) maintained its interaction with SGS3, while the LC2-deleted (U1-70k 1∼393aa) and LC1RmA variants did not (Fig. 6C). This result indicated that the C-terminal LC region indeed contributed to the U1-70k interaction with SGS3. Conversely, we tested whether the IDR of SGS3 contributes to this interaction. The C-terminal fragment of SGS3, which lacks an IDR, could still interact with U1-70k, suggesting that the IDR property of SGS3 is not essential for its interaction with U1-70k (Fig. 6H, S3I).

Given that the LC domain drives U1-70k self-oligomerization, and is also critical for its interaction with SGS3, we proposed the U1-70k multimerization driven by the LC domain promotes the U1-70k-SGS3 interaction (Fig. 6E). Importantly, the self-assembly of U1-70k was essential for its function in RNA silencing in vivo. This conclusion could be supported by the fact that LC1RmA mutated and LC domain deleted U1-70k could not rescue the LUC signal in *u1-70k-1* back to the level of E5-4 (Fig. 6F, G). These results further indicated that WT U1-70k could self-assemble into the condensate whereas the mutations of LC1 compromised this feature. Unexpectedly, we observed that U1-70k-LC1RmA even exhibited higher RNA binding activity than U1-70k but did not promote the transferring of RNA to SGS3 (Fig.6 I∼K, S5A, B, D). These results implied that the LC1 domain might have dual functions: on one hand, it could inhibit the accessibility of RRM of U1-70k to RNA. On the other hand, this very domain could promote LLPS and self-assembly, facilitating U1-70k/SGS3 interaction, which results in channeling ssRNA into SGS3 (Fig. S6D). If so, then LC1RmA might lead to greater exposure of the RRM and increased interaction with RNA, whereas the impaired multimerization of U1-70k LC1RmA may reduce SGS3 recruitment and ultimately reduce RNA loading onto SGS3 (Fig. S6D). These results suggested that the LLPS property of U1-70k could be critical for U1-70k function in PTGS.

### U1-70k forms co-condensate with SGS3

Given that the LC domain has LLPS property and its self-assembly is crucial for the U1-70k-SGS3 interaction, we hypothesized that U1-70k-SGS3 might form co-condensates. To test this, we first examined the LLPS feature of U1-70k and its variants (Fig. S7B). Recombinant mCherry-U1-70k indeed formed droplets at 1.5% PEG, while droplets disappeared upon the application of 10% 1,6-hexanediol (1, 6-HD) (Fig. 7A). Again, this biochemical feature was through the LC domain because U1-70k (235∼427aa), which only contains LC domain of U1-70k, behaved like WT U1-70k (Fig. 7A). Of note, U1-70kΔ5aa also acted as WT U1-70k, suggesting that the compromised function of *u1-70k-1* is irrelevant to its LLPS property (Fig. 7A). By contrast, U1-70k-LC1RmA could only form droplets at 2.25% PEG and the sizes of the droplets were smaller compared with other U1-70k variants (Fig. 7A, B). Furthermore, the condensates from mCherry-U1-70k-LC1RmA essentially disappeared by 10% of 1,6-HD treatment, whereas small droplets remained for WT U1-70k and other variants upon the 10% 1, 6-HD treatment (Fig. 7A). These results demonstrated that the LLPS property of U1-70k was weakened by R to A substitution in LC1 domain. Such results were reminiscent of mammalian U1-70k where basic-, acidic-amino acid repeated domain (BAD) in LC1 drives U1-70k into insoluble aggregates.^17^

**Fig. 7.**
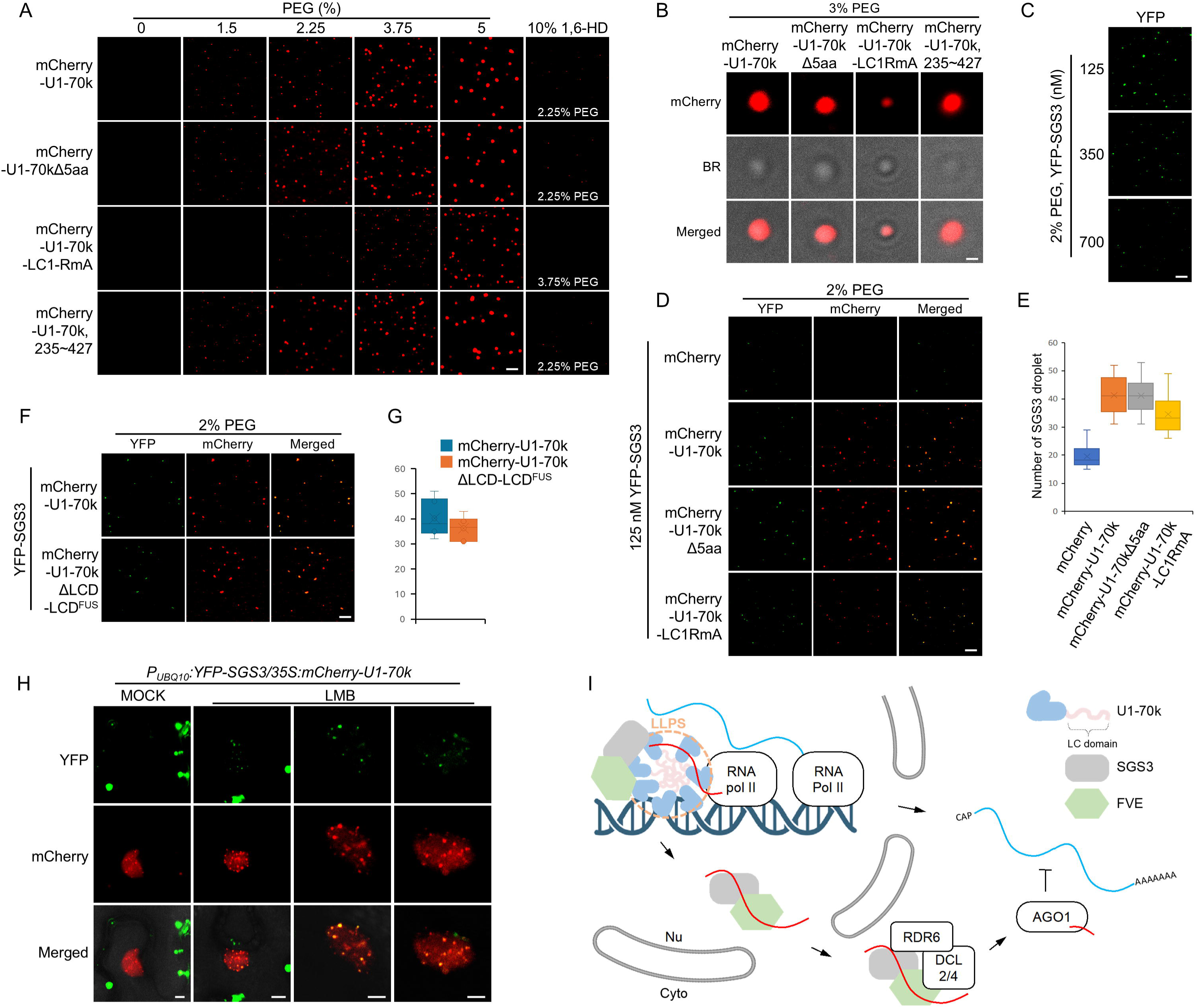
Liquid/liquid phase separation contributes to U1-70k/SGS3 interaction. (A, B) Confocal imaging shows that mCherry-U1-70k easily formed liquid droplets in vitro whereas this condensation was compromised by the LC1RmA mutation. 1 µM of mCherry tagged U1-70k, U1-70kΔ5aa, U1-70k-LC1-RmA and U1-70k (235∼427 aa) were used in the experiments. Scale bar, 5 µm (A) and 1 µm (B). (C) Confocal imaging shows that YFP-SGS3 begins to form condensates at a concentration of 125 nM in a 2% PEG solution. Scale bar, 5 µm. (D) Confocal imaging shows that U1-70k promotes the co-condensation of SGS3 via its LC domain. 1 µM of mCherry tagged U1-70k, U1-70kΔ5aa and, U1-70k-LC1-RmA were used in the experiments. Scale bar, 5 µm. (E) Statistic analysis of SGS3 droplet numbers in the panel (D) showed that LC1RmA mutation reduced the co-condensation activity of U1-70k. (F) Confocal imaging shows that the chimeric U1-70kΔLCD-LCD^FUS^ (deletion LC domain fused with LCD^FUS^) could form co-condensate with SGS3 in vitro. (G) Statistic analysis of SGS3 droplet numbers in panel (F). (H) Confocal imaging shows that SGS3/U1-70k co-condensates in the nucleus under LMB treated condition in tobacco leaf cells. Size bar, 5 µm. (I) A proposed model of U1-70k function in PTGS. U1-70k associates with the transgene locus and sequests nascent transcripts from Pol II co-transcriptionally. U1-70k could also form a liquid-like droplet and self-assembly driven by the C-term low complexity domain, which in turn recruits SGS3 and possibly FVE for relaying of the nascent RNA. The resultant ssRNA/SGS3/FVE is relocated from the nucleus to the cytosol to route onto RDR6 complex for dsRNA production prior to the action of DCL2/4 and downstream RISC activity.

SGS3 has been reported to show LLPS and this result was reproducible in our hands.^12^ Here, the minimum concentration was 125 nM for SGS3 condensation, and this concentration was further used as a sensitized condition for the co-condensation assay (Fig. 7C). Interestingly, upon addition of 1 uM of mCherry-U1-70k or -U1-70kΔ5aa, these proteins could indeed form co-condensates with SGS3 (Fig. 7D, E). Furthermore, the number of SGS3 condensates was doubled upon incubation of U1-70k or U1-70kΔ5aa, indicating that U1-70k could effectively recruit SGS3 into its condensates (Fig. 7D, E). By contrast, the application of mCherry-U1-70k-LC1-RmA could only elevate 70% more SGS3 condensates compared with the WT U1-70k (Fig. 7D, E). This result indicated that the LC domain of U1-70k plays a determining role in U1-70k/SGS3 co-condensates (Fig. 7D, E).

Next, we substituted LC domain of U1-70k with another IDR, the low-complexity domain from Fused in Sarcoma (LCD^FUS^), which efficiently undergoes LLPS in vivo and in vitro (Fig. S7B).^38,39^ mCherry-U1-70kΔLCD-LCD^FUS^ (LC domain deleted U1-70k fused with LCD^FUS^) promotes co-condensation of SGS3 with similar efficiency to the wild-type U1-70k protein, in terms of the number of SGS3 droplets (Fig. 7F, G). In fact, the IDR property of the LC domain, rather than its unique sequence content, such as BAD region, is crucial for the SGS3 interaction and co-condensation. Finally, we observed SGS3/U1-70k nuclear condensates in plant cells (Fig. 7H, S7B). These results indicate that U1-70k driven by its self-assembly and LLPS, pools SGS3 nearby the transgene loci and channels nascent RNA onto SGS3 and possibly FVE to embark on the PTGS tour.

## Discussion

How transgene transcripts are selected and enter the PTGS pathway has been a long-term mystery. Here, we deciphered this mystery, at least in part, and discovered that U1-70k, a well-known canonical component of spliceosome, routes nascent transcripts to the core components in RNA silencing pathway, SGS3 and FVE, which in return transport the transcripts from nucleus to cytosol and relay the substrates into RDR6 for dsRNA production. Several lines of evidence supported this notion: 1) our unbiased genetic screening recovered U1-70k as a new player in PTGS pathway, and its role is independent from its canonical function in U1 snRNP (Fig. 1, 2, 3); 2) multiple independent experiments validated U1-70k as a partner as FVE/SGS3 (Fig. 4); 3) both FVE and SGS3 can shuttle between the nucleus and the cytosol, creating opportunities to dynamically partner with U1-70k in the nucleus (Fig. 4); 4) U1-70k can bind to transgene loci and recruit the nascent transcripts from the loci in vivo (Fig. 5); 5) Although SGS3 does not seem to directly attach to chromatin, they can bind to nascent transcripts in WT plants (Fig. S4). Importantly, the association of SGS3 and FVE with transcripts replies on U1-70k (Fig. 5); and finally, 6) U1-70k could promote SGS3/FVE association with RNA in vitro in our reconstitution assays (Fig. 5). All these genetic and biochemical results prompted us to propose that U1-70k could directly request (or interact with) nascent transcripts from the transgene locus and promotes their loading of to SGS3 and FVE in the nucleus, which are subsequently translocated to cytosolic RDR6 for dsRNA synthesis (Fig. 5, 7I).

The fact that only nascent transcripts, but not mature mRNA, were recovered from SGS3 RIP in the nucleus strongly indicates that U1-70k-mediated relay of RNA substrate to SGS3 takes place co-transcriptionally and ahead of posttranscriptional processing (Fig. 5).

Mechanistically, U1-70k possesses some unique biochemical features, enabling its role in channeling RNA for silencing. First, U1-70k could directly interact with RNA Pol II, conferring itself a privilege to inherit the nascent transcripts from Pol II. Second, U1-70k has the RRM motif and can bind to RNA directly but with a relatively low affinity; this property warrants its ability to embrace transcripts for the first hand, while being able to relay transcripts to the next players in time. Third, U1-70k contains a unique LC region that not only enables U1-70k self-assembly but also displays LLPS (Fig. 6, 7). These properties provide a platform and condensate to enrich another IDP, SGS3, and possibly FVE. Finally, U1-70k does not typically leave from the chromatin or from at least not from nucleus, whereas SGS3/FVE could dynamically escape from the condensate due to a relatively weak association between SGS3/FVE and U1-70k (Fig. 7I). Thus, these advantages of U1-70k allow the protein to fetch nascent transcripts and transfer them directly to the downstream components in RNA silencing pathway.

It did not escape from our attention that some ta-siRNAs and phased siRNAs were also impacted but the others were not in loss-of-function mutants of *u1-70k-1*. Precisely, ta-siRNA1/2 and phased siRNAs, which are descendants from AGO1/miR173-RISC, are impacted by U1-70k; by contrast, ta-siRNA3, which is derived from AGO7/miR390 activity is not. Additionally, phased siRNAs from *Pentatricopeptide repeat (PPR)* genes and *AT3G23690*, which are processed by AGO1-miR161 and -miR393, respectively, were significantly reduced in *u1-70k-1* compared to WT. This contrast might result from the difference of subcellular compartmentations of the assembly of AGO1-miR173 and AGO7-miR390 RISCs. Although AGO1 mainly locates in the cytosol, AGO1-miRNA RISC is prevailingly believed to be assembled in the nucleus whiling acting in the cytoplasm.^40^ On the hand, AGO7-miR390 RISC is seemingly formed and functions exclusively in the cytoplasm. In this context, U1-70k might facilitate nascent *TAS* and *PPR* RNA substrates and contribute to the assembly of AGO1/miR173 or miR161/SGS3/*TAS1/2* RNA, promoting ta-siRNA and phased siRNA production. Alternatively, AGO7-miR390 might reply on a routine independent from U1-70k/SGS3 to match *TAS*3 substrates. Importantly, ta-siRNAs and phased siRNAs are critical weapons for host-induced gene silencing.^41,42^ It might be predicted that U1-70k might play dual roles in combating these pathogens by re-programming transcript splicing atlas in host while promoting the production of these host-induced gene silencing (HIGS).

A few other splicing regulators have also been reported to moonlight in RNA silencing. But their modes of action are distinct from the mechanism of U1-70k in promoting PTGS. PRP40, an auxiliary protein of U1 snRNP, and AAR2, an assembly factor of U5 snRNP, plant PRP24, and Brr2a, an RNA helicase in U4/U5/U6 snRNPs, regulates miRNA, but not siRNA, pathway in various ways.^43–46^ By comparison, PRP39a, another auxiliary protein of U1 snRNP, and SmD1, a core particle for U1/U2/U4/U5 snRNP, positively regulate PTGS.^47,48^ At a genetic level, these proteins act to inhibit nuclear RQC factors, accumulating aberrant transcripts as the templates for RDR6. However, their mechanisms are unclear at a biochemical level.

U1-70k is an evolutionarily conserved protein, and all contain LC domain that confers LLPS properties in a normal physiological condition and the aggregation under AD condition.^18–21^ Here, the U1-70k-mediated nuclear condensates function to relay the nascent RNA to the downstream component in PTGS. Whether additional components participate in the sorting of nascent transcripts into PTGS remains to be fully decoded. Similarly, it would be exciting to investigate whether U1-70k in other organisms share a similar noncanonical function to Arabidopsis counterpart in the future.

### Resource availability

#### Lead contact

Further information and requests for reagents should be directed to and will be fulfilled by the lead contact, Xiuren Zhang

#### Materials availability

All materials generated in this study are available from the lead contact upon request.

#### Data and code availability

The raw data produced during this study have been deposited into the GEO and Mendeley databases. They are publicly available as of the date of publication.

## Acknowledgments

This work was supported by grants from the NIH (R35GM151976) to X.Z.

## Author contributions

X.Z. conceived the project. T.O. and X.Z. designed the experiments. T.O. carried out most of functional and mechanistic studies for the manuscript. N.L., S.Z., and J.Z. provided some of genetic materials and intellectual input. K.L. initially identified *u1-70k-1* via the genetic screening and Y2H screening of U1-70k’s partners. C.L. performed bioinformatical analysis. S.L. provided *luc7a/b/rl* triple KO mutant seeds. T.O. drafted the manuscript, and X.Z. thoroughly edited the draft.

## Material and Methods

### Vector construction

To obtain pK-pU1-70k-FM-gU1-70k, *U1-70k* promoter, a fragment of 1500 bp upstream of the start codon ATG of *U1-70k* gene, was amplified using Thermococcus kodakaraenis (KOD) DNA polymerase (Novagen). The *U1-70k* promoter and the XbaI-linearized pK-MCS-FM-DC vector were mixed with HiFi DNA Assembly Master Mix (HiFi enzyme, NEB) to generate pK-pU1-70k-FM-DC followed by DNA sequencing. Genomic DNA of *U1-70k* (*gU1-70k*) was amplified and cloned into pENTR vector by TOPO reaction and validated via Sanger sequencing. The *gU1-70k* was switched from pENTR-gU1-70k to pK-pU1-70k-FM-DC via a LR Clonase II enzyme reaction (Gateway system, Thermo fisher).

For the pK-pRDR6-FM-DC construction, pK-MCS-FM-DC was cut by Xba I which digest prior to the FM tag region. The promoter region of *RDR6* was amplified by KOD DNA polymerase and the PCR product was introduced into Xba I cut pK-MCS-FM-DC via HiFi reaction.

The CDS clone of *U1-70k* and *SGS3*, and the series of truncations used for plants transformation and Y2H were cloned by using T4 ligations or NEBuilder HiFi DNA Assembly Master Mix and the Gateway system. pENTR-MCS vector was cut by Sal I/Xba I and the DNA fragment of interests were amplified by KOD DNA polymerase with the sets of primer which contains Sal I and Xba I restriction enzyme site in the 5’- and 3’-primer, respectively. The PCR products were digested by Sal I/Xba I and the fragments were ligated into pENTR-MCS by T4 DNA ligase (NEB) or introduced into pENTR-MCS by using HiFi reaction. The recombinant DNA, pENTR-MCS containing the gene of interest, was subjected to LR recombination with a specific destination vector, based on the purpose of the cloning, such as plant transformation or Y2H assay. pK-35S-FM-DC, pK-pRDR6-FM-DC, pH7GWY2-35S-YFP-DC, pBA-35S-mCherry-DC, pH-pUBQ10-YFP-DC, pBA-nYFP-DC, pBA-cYFP-DC and pBA-pRDR6-YFP-DC were used for transformation of plants and pGBK-T7-GW and pGAD-T7-GW were used for the Y2H assay. The pENTR-MSC clone was digested with EcoR V or Apa I for recombination into pGBK-T7-GW, as both vectors are resistant to kanamycin.

For the cloning of *u1-70k-1* CDS, which is 15 nt deleted form of *U1-70k* identified in the *ars 2-1* mutant, U1-70k was amplified through PCR by using cDNA obtained from *u1-70k-1* mutants. The PCR product was cloned into pENTR vector via D-topo reaction.

For cloning of the U1-70k-LC1RmA, all the Arginine in LC1 region, about 281 nt long, was substituted to Alanine by overlapping PCR reaction with three 110 bases long primer which covers all LC1 region and contains the mutated codon that generate Alanine instead of Arginine. Prior to the PCR reaction, the primers were loaded into urea-PAGE. The top bands were eluted and used in the PCR reaction. The amplified DNA fragment was cloned into the destination vectors through the process as described above.

For cloning of artificial miRNA U1-70k (amiR-U1-70k) and amiR-U1C constructs, a template of pENTR-amiR159 was amplified with a set of primers that contain the amiRNA sequences complementary to the U1-70k or U1C mRNA. The resultant PCR products were cut by Xma I/Bgl II, and then ligated into the Xma I/BgI II-digested pENTR/D-amiR backbone to generate pENTR/D-amiR-U1-70k or -U1C, accordingly. The amiR-U1-70k and -U1C were cloned into pK-35S-DC via LR recombination.

For the protein expression clones for EMSA assays, pET28-HIS-SUMO-HYL vector was digested by BamH I and Sal I results in cutting out of HYL1 CDS. *U1-70k*, *u1-70k-1* and *SGS3* CDS were amplified by PCR and the HiFi builder DNA assembly system was used for the subsequent cloning.

For the in vitro LLPS assay, pET28-HIS-SUMO-HYL1 vector was modified to pET28-HIS-YFP-HYL1 and pET28-HIS-mCherry-HYL1. pET28-HIS-SUMO-HYL1 was cut by Nde I/BamH I to cut out the SUMO tag and amplified YFP and mCherry were fused with the Nde I/BamH I digested pET28-HIS-HYL1 via HiFi reaction. The pET28-HIS-YFP-HYL1 and pET28-HIS-mCherry-HYL1 were cut by BamH I and Sal I. The amplified SGS3 CDS was cloned into pET28-HIS-YFP- and U1-70k, U1-70kΔ5aa, U1-70k-LC1-RmA and U1-70k 235∼427 were introduced into pET-HIS-mCherry-via HiFi reaction. To obtain the pET28-HIS-mCherry-U1-70kΔLCD-LCD^FUS^, U1-70k N-term region about 1 to 720 nt and the LCD^FUS^ were amplified via PCR. These two fragments were fused with BamH I/Sal I cut pET-HIS-mCherry-via HiFi reaction.

### Plant materials and growth conditions

Plants were grown on soil in 16-hour light/8-hour dark or MS plates in 12-hour light/12-hour dark at 22°C. *A. thaliana* ecotype Landsberg (Ler), Columbia (Col-0), E5-4 (Ler; PAGO10-GFP-PHB-LUC), were used for this study. Ec5-4, E5-4 crossed with Col-0 for seven times, was crossed with *u1a* (SALK_074230C) and *luc7a/b/rl* to generate Ec5-4/*u1a* and Ec5-4/*luc7a/b/rl,* respectively. Homozygous of *u1a* and *luc7a/b/rl* mutation in Ec5-4 were confirmed by genomic DNA PCR using primers indicated in table S1 in the F2 generation. EMS mutagenesis, mutant screen and LUC assays for identifying *ars2-1* were performed as described previously.^11^ *u1-70k-1* was crossed with L1-GUS and *35S:CHS* lines for generating L1-GUS/*u1-70k-1* and *35S:CHS/u1-70k-1,* respectively, and homozygous of each genetic lines were identified by PCR in F2 generation.

### RNA-seq and sRNA-seq analysis

TRIzol was used to extract total RNA from 7-day-old seedlings grown on MS plates. mRNA and sRNA sequencing library constructions and bioinformatic analysis were performed as described.^33,49^ Illumina sequencing was performed by the Novogene, US.

For the differential splicing analysis, the trimmed reads were mapped to the Arabidopsis genome (v.TAIR10) with HISAT2 (v.2.2.1). The alignment results were converted to BAM format, indexed by SAMtools (v.1.9). Differentially spliced transcripts were identified by rMATS (v.4.1.2).

### U1-70k antibody generation

To avoid targeting the conserved RRM domain, which is common among RNA-binding proteins in Arabidopsis, we used the N-terminal and C-terminal regions of the U1-70k CDS for epitope selection in antibody production. pET28-HIS-SUNO-MCS was cut with BamHI/SalI and 1∼453 nt (1∼151 amino acids, N-epitope) with a stop codon, and 733∼1284 nt (245∼427 amino acids, C-epitope) were cloned into the vector by HiFi DNA assembly (NEB) reaction. The recombinant plasmids were transformed into BL21DE3 and the recombinant proteins were induced by adding 0.5 mM IPTG at the optical density at 600 (OD_600_) 0.8 grown at 18°C for 16 hours. The epitope induced cells were dissolved in Lysis buffer (20 mM Tris-HCl pH 8.5, 500 mM NaCl, 2% glycerol, 0.1% Triton X-100, 25 mM imidazole, 1 mM PMSF) and Lysis buffer 2 (20 mM Tris-HCl pH 8.5, 1 M NaCl, 2% glycerol, 0.1% Triton X-100, 25 mM imidazole, 1 mM phenylmethylsulfonyl fluoride (PMSF)), respectively, and lysed by using LM20 microfluidizer with 25,000 psi for three times. The lysates were centrifuged with 20,000 RPM at 4°C for 30 min and subjected to affinity purification using HIS-TRAP HP 5 ml column on Akta-pure FPLC system (GE). The Elutes were dialyzed in 0.01% HIS-SUMOPROTEASE (v/v), 20 mM Tris-HCl pH 8.0, 400 mM NaCl, 1 mM β-mercaptoethanol at 4°C for 16 hours and subsequently incubated with Ni-NTA beads for 1 hour, 4°C to remove those HIS-SUMOPROTEASE and HIS-SUMO proteins from the elutes. N-epitope was concentrated with Amicon Ultra Centrifugal Filter, 10 kDa MWCO column (Millipore sigma) and C-epitope was subjected to size exclusion chromatograph with the buffer contains 20 mM Tris-HCl pH 8.0, 1M NaCl, 320 mM imidazole, by using Hiload 16-600 superdex 75 pg column on an AKTA-pure FPLC system (GE). The collected fractions were dialyzed and concentrated with the same buffer and method used for N-epitope preparation. The concentration of purified epitopes was adjusted to 2 ug/ul in the mixed epitope solution and sent out for antibody production to Rockland, USA (Fig. S1G). U1-70k antibody was purified from the rabbit total blood serum by using protein A magnetic beads (Thermo Fisher). The U1-70k antibody was validated by using E5-4, *u1-70k-1*, E5-4/*U1-70k KD-2*, and *35S:FM-U1-70k* (Fig. S1H).

### Western blot

SDS-PAGE, PVDF membrane transfer and western blot assays were performed as described previously.^50^ anti-Myc (GenScript, A00704), anti-actin (Sigma-Aldrich, A0480), anti-UGPase (Agrisera, AS14 2813), anti-histone 3 (Agrisera, AS10 710), anti-SGS3 (Agrisera, AS15 3099), anti-HA (Sigma-Aldrich, H9658), and anti-U1-70k (homemade) were used for the first-round western blot. A goat-developed anti-rabbit-HRP (GE Healthcare, catalog no. NA934) and anti-mouse immunoglobulin G-HRP (GE Healthcare, catalog no. NA931) were used for second round western blot. The membranes were subsequently treated with ECL solution and ChemiDoc MP imaging system, Biorad was used for the signal detection.

### Nuclear-cytoplasmic fractionation and Nuclear-cytoplasmic RNA extraction

0.3 g of two-week-old seedlings were homogenized with liquid nitrogen into a fine powder and dissolved into 3 ml of lysis buffer (20 mM Tris-HCl pH 7.5, 20 mM KCl, 2.5 mM MgCl_2_, 25% glycerol, 250 mM sucrose, 5 mM dithiothreitol (DTT), 20 μM MG-132, 0.5× protease inhibitor (Roche), 100 U ml^−1^ SUPERase-In RNase Inhibitor). The lysate was thoroughly mixed with gentle inverting and filtered through two layers of Miracloth twice. 500 μl of lysate was sampled as a total fraction and rest of the extracts were centrifuged at 1,000g for 10 min at 4 °C. 1 ml of the supernatant was collected as cytoplasmic fraction, followed by another centrifuged at 13,000g for 10 min at 4°C. The pellet was washed with 5 ml of nuclear washing buffer twice (20 mM Tris-HCl pH 7.5, 20 mM KCl, 2.5 mM MgCl2, 25% glycerol, 250 mM sucrose, 0.2% Triton X-100, 100 U ml^−1^ SUPERase-In Rnase Inhibitor). After washing, the pellet was directly dissolved into Xeno-sepa-TR solution (Xenohelix) for RNA extraction or resuspended with 1 ml of nuclear washing buffer for western blot. Each fraction was mixed with same volume of SDS sample buffer and subsequently loaded SDS-PAGE for western blot. RNA extraction from total and cytosolic fraction was performed by using total RNA extraction kit (Xenohelix).

### Leptomycin B treatment

Seven-day-old seedlings grown on the mesh-covered MS plate were transferred to 3 ml liquid MS medium for 12 h, then 2.5 μM leptomycin B (Enzo Life Sciences) or an equal amount of 100% ethanol was added to the medium, which served as a mock. The samples were subjected to vacuum with −25 inHg for 5 min three times and incubated in the light chamber for 6 hours. For the tobacco leaves, 1 μM leptomycin B (Enzo Life Sciences) or 0.54% of Ethanol were infiltrated 2 days after agrobacterium infiltration and incubated for 6 hours.

### Coimmunoprecipitation

Two-week-old seedling were ground into fine power in the liquid nitrogen. 0.2 g Powder was mixed in 0.6 ml of IP buffer (40 mM Tris-HCl pH 7.4, 150 mM KCl, 5 mM MgCl2, 0.5% Triton X-100, 1% glycerol, 5 mM dithiothreitol (DTT), 20 μM MG-132, 0.5× protease inhibitor (Roche)). The total protein extracts were centrifuged at 13,000 g for 10 min at 4°C twice. 40 μl of extracts were sampled as an input and anti-c-Myc-magnetic beads (Pierce) or SGS3 antibody with protein A magnetic beads (20nvitrogen Dynabeads) were added to 400 μl of the extracts and incubated on the rotator at 4°C for one hours. Beads were briefly washed three times with wash buffer (40 mM Tris-HCl pH 7.4, 150 mM KCl, 5 mM MgCl_2_, 0.2% Triton X-100, 1% glycerol). 40 μl of 1x SDS sample buffer was mixed with beads and 10 μl of 6x SDS sample buffer was added to the input sample. After boiling, 20 μl of each sample were loaded into SDS-PAGE for western blot.

For the nuclear fraction CoIP, 0.8 g of plants powder in 8 ml of nuclear fractionation lysis buffer was used for nuclear fractionation. The isolated nuclear pellet was mixed with 0.6 ml of IP buffer, and the same process described above was used for the subsequent steps.

### Ribonucleoprotein Immunoprecipitation-polymerase chain reaction

2.5-week-old seedlings were submerged in cross-linking buffer (20 mM Tris-HCl pH 7.4, 1 mM PMSF, 1 mM EDTA, 0.4 M sucrose and 1% formaldehyde) and vacuumed four times by −25 inHg for 5 min. The cross-linked plants samples were vacuumed three times with de-cross-linking buffer (20 mM Tris-HCl pH 7.4, 0.4 M sucrose and 125 mM Glycine) and then washed with DEPC water for three times. The nuclei pellet was prepared from 0.8 g of grounded sample powder and was thoroughly dissolved into 300 μl lysis buffer (40 mM Tris-HCl pH 7.4, 500 mM KCl, 5 mM MgCl_2_, 1% SDS, 1 mM DTT, 1% Triton X-100, 2% glycerol, 20 μM MG-132, 0.5x protease inhibitor and 100 U ml^−1^ SUPERase-In Rnase Inhibitor). The extracts were centrifuged at 4°C, 13,000 g for 10 min and 120 μl of the supernatant was sampled for input. 150 μl of nuclear lysate was diluted with 600 μl of dilution buffer (40 mM Tris-HCl pH 7.4, 100 mM KCl, 5 mM MgCl2, 1 mM DTT, 0.2% Triton X-100, 2% glycerol, 20 μM MG-132, 0.5x protease inhibitor and 100 U ml^−1^ SUPERase-In Rnase Inhibitor) followed by co-incubation with anti-c-Myc-magnetic beads (Pierce) or SGS3 antibody with protein A magnetic beads (Pierce) at 4°C for 1 hours. The beads were washed for three times with washing buffer 1 (40 mM Tris-HCl pH 7.4, 200 mM KCl, 5 mM MgCl_2_, 0.5% Triton X-100, 2% glycerol) and two times washed with washing buffer 2 (40 mM Tris-HCl pH 7.4, 500 mM NaCl, 5 mM MgCl2, 0.2% Triton X-100, 2% glycerol). The beads were resuspended with 125 μl of RiP lysis buffer and 25 μl of slurry was sampled for western blot. 100 μl of IP and input samples were mixed with proteinase K solution (40 mM Tris-HCl pH 7.4, 200 mM NaCl, 10 mM EDTA, 1% SDS, 0.4 mg proteinase K) and incubated at 65°C with gentle shaking for 16 hours. RNA was extracted from the sample by using TRIzol method and followed by Dnase treatment, reverse transcription and PCR for analysis. 5 μl of inputs and the 25 μl of IP products were used for western blot.

### Chromatin immunoprecipitation–polymerase chain reaction

ChIP-PCR assay was performed as described.^51^ 2.5 weeks old plants were used in the ChIP-PCR and IP was performed with an anti-Myc antibody (Sigma-Aldrich, C3956). PCR primers are listed in table S1.

### Cellular localization assay and BiFC assay

Protoplast transfection and tobacco infiltration were performed as described.^52^ Leaves of four weeks old Col-0 grown on soil in 10-hour light/14-hour dark were used for the protoplast preparation and 3.5 weeks old tobacco leaves were used for the infiltration. The fluorescence signal from the protoplasts and tobacco leaves were visualized by Leica stellaris 5 laser-scanning confocal microscope.

### GUS staining

GUS staining was performed as described previously.^2^ Five-day-old seedlings of L1-GUS and L1-GUS/*u1-70k-1* F3 were used for the GUS staining.

### Y2H assays

Yeast transformation was performed according to the manual of Gold Yeast Two-Hybrid System by the manufacturer (Clontech). After the transformation, half of transformed yeast cells were spread on -Trp/-Leu (2 out) media and the rest half were spread on -Trp/-Leu/-His (3 out) plate. After three days of incubation, colonies were selected from the 3 out plates, in case cells had grown on the media. If no growth was observed, colonies were collected from the 2 out plates and transferred to liquid culture in YPDA medium for overnight incubation. Next day, the density of cells in the culture was synchronized to OD_600_ = 0.3 and spotted on to 2 out and -Trp/-Leu/-His/-Ade (4 out) plate with 10 times dilution. The colonies from each plate were observed after three days of incubation.

### Expression and purification of recombinant proteins

6×His-SUMO-U1-70k, 6×His-SUMO-U1-70kΔ5aa, 6×His-SUMO-U1-70k-LC1RmA, 6×His-SUMO-SGS3, 6×His-mCherry, 6×His-mCherry-U1-70k, 6×His-mCherry-U1-70kΔ5aa, 6×His-mCherry-U1-70k-LC1RmA, 6×His-mCherry-U1-70k 237∼427, 6×His-mCherry-U1-70kΔLCD-LCD^FUS^ and 6×His-YFP-SGS3 were expressed in *E. coli* BL21 (DE3) cells. The transformed BL21 cells were grown in LB at 37°C until the OD_600_ reaches to 0.6, then the cells were cooled down on ice for 15 min. 0.4 mM isopropyl-β-d-thiogalactoside was added to the flask and the cells were grown at 16°C for 16 hours.

For purification of 6×His-SUMO-U1-70k, 6×His-SUMO-U1-70kΔ5aa, 6×His-SUMO-U1-70k-LC1RmA, 6×His-mCherry-U1-70k, 6×His-mCherry-U1-70kΔ5aa, 6×His-mCherry-U1-70k-LC1RmA and 6×His-mCherry-U1-70k 237∼427, the harvested cells from 2 to 4 L culture were thoroughly resuspended into lysis buffer (20 mM Tris-HCl pH 7.4, 1 M NaCl, 2% glycerol, 0.1% triton X-100, 25 mM imidazole, 1 mM PMSF). The cells were lysed by using LM20 Microfluidizer at 25,000 psi for three times on ice. The lysates were centrifuged at 20,000 rpm for 20 min at 4°C. The supernatant was filtered through 0.2-μm membrane filter, and the extracts were loaded into a HisTrap HP column (GE Healthcare) by using Akta-pure FPLC system. The column was washed with washing buffer (20 mM Tris-HCl, pH 7.4, 1 M NaCl, 2% glycerol, 1 mM β-mercaptoethanol) while gradually increasing the imidazole concentration. The washing steps included: 30 ml of 32 mM imidazole, followed by 30 ml of 45 mM imidazole, 30 ml of 60 mM imidazole, and finally, a linear gradient from 60 mM to 120 mM imidazole over 25 ml. Then the recombinant protein was eluted by adding 320 mM of imidazole in washing buffer. This washing/elution method was developed to obtain highly concentrated 6xHIS-SUMO-U1-70k fractions, as purified U1-70k tends to bind to the inner membrane of the Amicon concentration column and cannot be concentrated efficiently. Eluted fractions were collected based on the UV absorbance signal at 280 nm and were subsequently subjected to SDS-PAGE to assess protein quantity. 3 ml of purified recombinant protein from selected fractions were loaded into Hiload 16/600 superdex 200 pg column with the buffer contains Tris-HCl pH 7.4, 1M NaCl, 400 mM imidazole. The fractions were sampled based on the UV signal and the quantity and quality of recombinant protein was confirmed by SDS-PAGE. The highly concentrated fractions with fine quality were dialyzed with dialysis buffer 1 (Tris-HCl pH 7.4, 500 mM NaCl, 25% glycerol, 1 mM β-mercaptoethanol) for 6xHIS-SUMO tagged protein. 6xHIS-mCherry tagged proteins were dialyzed in dialysis buffer 2 which contains 5% glycerol instead of 25%. The dialyzed recombinant proteins were stored in −80°C.

6×His-SUMO-SGS3, 6×His-mCherry, 6×His-mCherry-U1-70kΔLCD-LCD^FUS^, and 6×His-YFP-SGS3 were purified following a similar protocol as described above, with modifications in buffer pH and salt concentration for each purification. 6×His-SUMO-SGS3, 6×His-mCherry, and 6×His-YFP-SGS3 were purified using lysis buffer (20 mM Tris-HCl pH 8.5, 500 mM NaCl, 2% glycerol, 0.1% triton X-100, 25 mM imidazole, 1 mM PMSF), washing buffer (20 mM Tris-HCl, pH 8.5, 500 mM NaCl, 2% glycerol, 1 mM β-mercaptoethanol), gel filtration buffer (20 mM Tris-HCl pH 8.5, 500 mM NaCl). 6×His-mCherry-U1-70kΔLCD^FUS^ was purified with lysis buffer (40 mM Tris-HCl pH 8.0, 800 mM NaCl, 2% glycerol, 0.1% triton X-100, 25 mM imidazole, 1 mM PMSF), washing buffer (40 mM Tris-HCl, pH 8.0, 800 mM NaCl, 2% glycerol, 1 mM β-mercaptoethanol), gel filtration buffer (40 mM Tris-HCl pH 8.0, 800 mM NaCl). The same dialysis buffer was used for these proteins.

### In vitro transcription and labeling of RNA

For in vitro transcription, the forward strand of minimal T7 promoter and reverse strands of the template, which contains T7 promoter and reverse-complement sequence of the 38-nt ssRNA, was generated and further purified through urea-PAGE method as demonstrated above. 10 pmol of each strand was mixed in 30μl of 10 mM Tris-HCl pH 8.0 solution and placed on the heat block with 95°C. Then the heat block was placed in the room temperature for 3 hours to achieve dimerization of DNA and then T7 in vitro transcription was performed to obtain 38-nt ssRNA as described previously.^53^ For Cy5 labeling, 1 mM of Cy5-UTP was added into reaction mix with 1 mM of non-labeled UTP during T7 transcription. 5′ end labeling of ssRNA were performed as described ^7^.

### Electrophoretic mobility shift assays

Different concentration of recombinant proteins was prepared in with buffer composition, 20 mM Tris-HCl pH 7.4, 25% glycerol, 500 mM NaCl. 4 μl of recombinant protein was added into 20 μl total EMSA mixture (21 mM Tris-HCl pH 7.4, 1 mM MgCl_2_, 0.5 mM DTT, 5 U ml^−1^ SUPERase-In Rnase Inhibitor, 0.15 μM IZ-^32^P-ssRNA) and incubated on ice for 20 min, then the samples were mixed with 6x DNA loading dye, then loaded into 1.8% agarose gel. After migration, agarose gel was incubated with fixation buffer (40% ethanol, 10% acetic acid, and 5% glycerol) with gentle agitation for 30 min. Then the gel was dried overnight, and the radioisotope signal was analyzed by typhoon FLA9500 phosphor-imager.

### In vitro liquid-liquid phase separation assay

Prior to the in vitro condensation assay, all protein samples were centrifuged at 13,000g for 10 minutes, and the supernatants were transferred to new tubes to remove aggregates. The recombinant protein was diluted into final concentration of 20 mM Tris-HCl pH 7.4, 100 mM NaCl, 1% glycerol and diverse concentration of the crowding reagent, poly-ethylene glycol 4000.

To disrupt liquid like droplet formation, 10% of 1,6-hexanediol was added into reaction mix. The condensates were captured by Leica stellaris 5 laser-scanning confocal microscope with a HC PL APO 63×/1.40 oil-immersion objective and LAS X Life Science Microscope software.

---

**Fig. S1.** Identification of *u1-70k-1* mutant via a genetic screening. (A) The reporter gene structure in the *E5-4* parent line. (B) Next-generation mapping (NGM) analysis identified the causative loci in *ars2-1* mutant from the F2 population. (C) NGM analysis pinpointed two candidate genes for *ars2-1* in 1.8 Mb of chromosome 3. (D) Complementation assays of LUC signal result shows that *At3g50670* (*U1-70k)* gene is causative for the *ars2-1* mutation. Scale bars, 5 mm. (E) A schematic diagram of the domain structure of U1-70k. Computational modeling suggested that two unique epitopes in U1-70k are labeled at the bottom. (F) Phylogenetic analysis shows that that RRM domain of U1-70k is highly conserved across animals and plants. (G) SDS-PAGE result demonstrates the quantity and quality of purified immunogens used in U1-70k antibody production. (H) Western blot assay indicates that the home-made antibody could detect the endogenously expressed U1-70k protein under the native or constitutive promoter. Total crude extracts from rosette leaves of three-week-old plants of the indicated genetic lines were used in Western blot assay. Cross-reactive bands were marked with red asterisks. Be noted that the U1-70k was not detectable in the *U1-70k* KD-2 lines. (I) Design of amiRNAs that specifically target the *U1-70k* transcripts. (J) *U1-70k* KD plants showed severe defects in growth and development. 5-week-old plants (top) and 3.5-week-old plants (bottom).

**Fig. S2.** High reproducibility of sRNA-seq and RNA-seq in assessing the disruption of the PTGS pathway in selected mutant vs WT plants. (A) Principal component analysis (PCA) of sRNA-sequencing shows a well-clustered pattern for biological replicates. (B) Northen blot assay showed that the accumulation of selected founding miRNAs was comparable between *u1-70k-1* and E5-4. U6 snRNA serves as a loading control. (C) qRT-PCR assay demonstrated the expression levels of *AGO10* and miR165/166 target genes were comparable in *u1-70k-*1 and E5-4. (D) The PCA analysis demonstrates clear clustering of biological replicates of each mutant or WT plants in RNA sequencing. (E) PCA plot analysis further validated the clustered characteristic of *u1-70k-1* replicates vs E5-4. (F) Computational analysis shows that the phased siRNA from PPR genes processed by AGO1-miR161 is reduced in *u1-70k-1* vs WT. The asterisks indicate *p*-value from student’s *t*-test (***, *p* < 0.001; **, *p* < 0.005; *, *p* < 0.02). (G) Design of amiRNAs that specifically target the *U1C* transcripts. (H) RT-PCR assay showed that reduced expression level of *U1C* in *U1C* KD lines vs WT. *UBC10* was used for an internal control and the band intensity was measured by image J. (I) RNA-seq did not reveal the significant changes of the expression levels of RNA quality control genes and sRNA pathway genes in *u1-70k-1* vs E5-4. The y-axis indicates the normalized expression level presented as the mean of three replicates of normalized reads.

**Fig. S3.** Additional experimental replicates support the U1-70k/SGS3 interaction in the nucleus. (A) Y2H screening showed that U1-70k interacts with SGS3 but not with other components in RNA silencing exemplified by DCL2, DCL3, DCL4, RDR2, or RDR6 in yeast. (B) Negative controls for BiFC assay in the panel (Fig. 4C). (C) BiFC assay using Arabidopsis protoplast independently validated that either U1-70k and U1-70kΔ5aa could interact with SGS3. (D) YFP-SGS3 and mCherry-U1-70k were expressed in Arabidopsis protoplast and visualized in Leica stellaris 5 laser-scanning confocal microscope. The arrowheads indicate the locations of the nucleus. (E) Western blot assays of additional biological replicates exhibited the clear accumulation of SGS3 in the nucleus upon the LMB treatment. The involved proteins were detected with anti-bodies against endogenous proteins, but RDR6 signal was relatively weaker. (F) Western blot analysis of nuclear/cytosol fractionation assay using an anti-Myc antibody showed that FM-RDR6 was highly distributed in the cytosol and barely visible in the nucleus. UGPase and H3 served as positive controls for the cytosol and nucleus fractions (E, F). (G) Y2H assay showed that the five-aa deletion of U1-70k (U1-70kΔ5aa) did not affect its interaction with SGS3 and FVE. (H, I) Y2H assay did not show auto-activity of indicated proteins.

**Fig. S4.** Additional datasets independently validated the quality of nuclear/cytosol fraction for RT-PCR and chromatin/RNA association of U1-70k, SGS3 and FVE. (A) Western blot assay validated the clear separation of nuclear/cytosol fractions from E5-4 and *u1-70k-1* mutant used in Fig. 5B and S4C. (B) An agarose gel image showed the quality of RNA in total/nuclear/cytosolic fraction used in Fig. 5B and S4C. (C) RT-PCR assays using two sets of primers showed the distribution of nascent and processed E5-4 transcripts in the nuclear and cytosolic fractions. Be noted that the silencing of *GFP-PHB-LUC* transcript in E5-4 predominantly occurred in the cytosol. *AtUBC10* was used as an internal control and red box with dashed line indicates the data used in Fig. 5B. The asterisk and arrowhead indicate nascent *GFP-PHB-LUC* transcript and matured transcript, respectively. (D) A schematic diagram showed the locations of primer sets used in ChIP/RIP-PCR in the *E5-4* reporter construct in this figure. (E) Three independent replicates of ChIP-PCR assays exhibited that SGS3 did not directly associate with the *GFP-PHB-LUC* locus, either in the E5-4 parent line or *u1-70k-1*. (F) RIP-PCR assay using total crude extract demonstrated that SGS3 bound to *GFP-PHB-LUC* transcripts in a U1-70k-dependent manner. Be noted: the transgene *SGS3* is controlled by the *RDR6* promoter, as used by Blagojevic et al., as the endogenous regulatory element of SGS3 is complicated and will be studied elsewhere.^54^ (G) RIP-PCR results showed that *GUS* transcript was recovered from SGS3 IP in the WT plants but not in *u1-70k-1*.

**Fig. S5.** Additional replicates of EMSA showed ssRNA interaction with U1-70k/SGS3/FVE. (A) SDS-PAGE showed the purified recombinant protein used in EMSA assay (B) U1-70k, U1-70kΔ5aa, U1-70k-LC1RmA, and SGS3 showed similar binding pattern but with different affinities with homogenous ssRNA consists with 3 guanines and 44 adenines (G3A44) in the EMSA assay. (C) Additional biological replicates of EMSA indicate dramatically reduced RNA binding affinity of U1-70kΔ5aa compared with wild-type U1-70k (Fig. 5G). These results were used to calculate the *K*_d_ in Fig. 5J. (D) Repeated EMSA results of U1-70k, U1-70k-LC1RmA, SGS3, and FVE were used in calculating *K*_d_ in Fig. 5 via imaging J following the manufacturer’s instruction. (E) EMSA assay demonstrated that U1-70k, but U1-70kΔ5aa, promoted G3A44 loading into SGS3.

**Fig. S6.** Computational analysis revealed that U1-70k contained a lengthy and conserved IDR that potentiates the self-assembly and its interaction with SGS3. (A) Computational analysis predicted the presence of a short and a lengthy IDR in the N-terminal and C-terminal regions, of U1-70k, respectively (www.pondr.com). Boxes with blue dashed line indicate N-terminal IDR (N-IDR) and low-complexity region. (B) Amino acid sequence alignment indicated that the N-terminal region of U1-70k was well conserved in plants but not in mammals. (C) LC domain of U1-70k is well conserved across diverse plant and mammal species. (D) A hypothetical model demonstrates enhanced ssRNA binding affinity of U1-70k-LC1RmA and reduced ssRNA binding to SGS3 in U1-70k-LC1RmA.

**Fig. S7.** Confocal microscope imaging shows that U1-70k could form co-condensates with SGS3 in the nucleus under LMB treated condition in plant cells. (A) SDS-PAGE showed the purified recombinant protein used in vitro LLPS assay. (B) YFP-SGS3 and mCherry-U1-70k were expressed in *N. Benthamiana* leaves and visualized in Leica stellaris 5 laser-scanning confocal microscope. The arrowheads indicate the locations of the nucleus. Be noted: the overlapping of the speckles was indicated by the yellow foci in the nucleus in the merged panels.

