## Supplementary figures for "The spliceosome component U1-70k loads nascent transcripts onto FVE/SGS3 via co-condensation to embark on RNA silencing"

Fig. S1

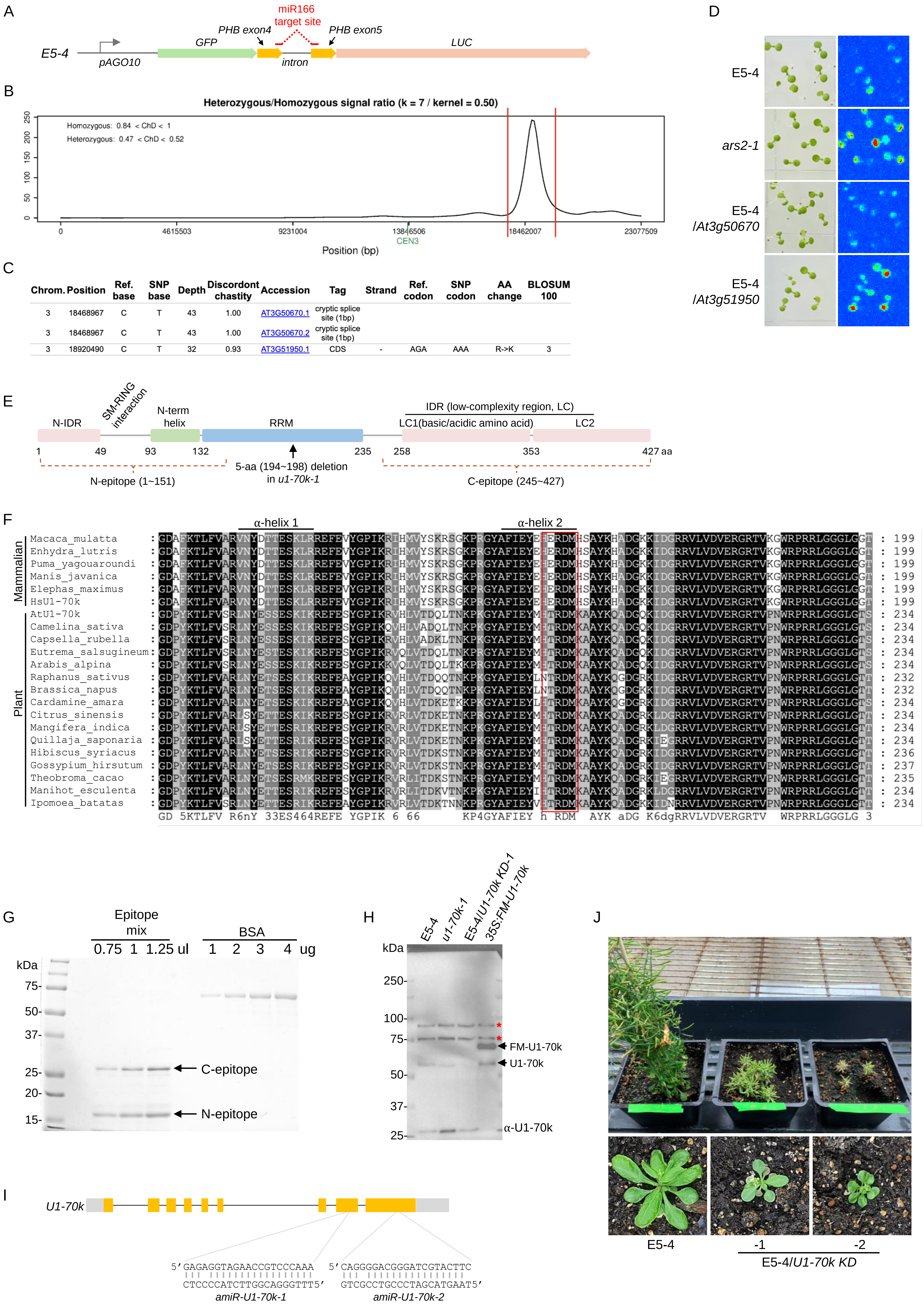


| Chrom. | Position | Ref. base | SNP base | Depth | Discordont chastity | Accession | Tag | Strand | Ref. codon | SNP codon | AA change | BLOSUM 100 |
|---|---|---|---|---|---|---|---|---|---|---|---|---|
| 3 | 18468967 | C | T | 43 | 1.00 | AT3G50670.1 | cryptic splice site (1bp) | | | | | |
| 3 | 18468967 | C | T | 43 | 1.00 | AT3G50670.2 | cryptic splice site (1bp) | | | | | |
| 3 | 18920490 | C | T | 32 | 0.93 | AT3G51950.1 | CDS | - | AGA | AAA | R->K | 3 |

Fig. S2

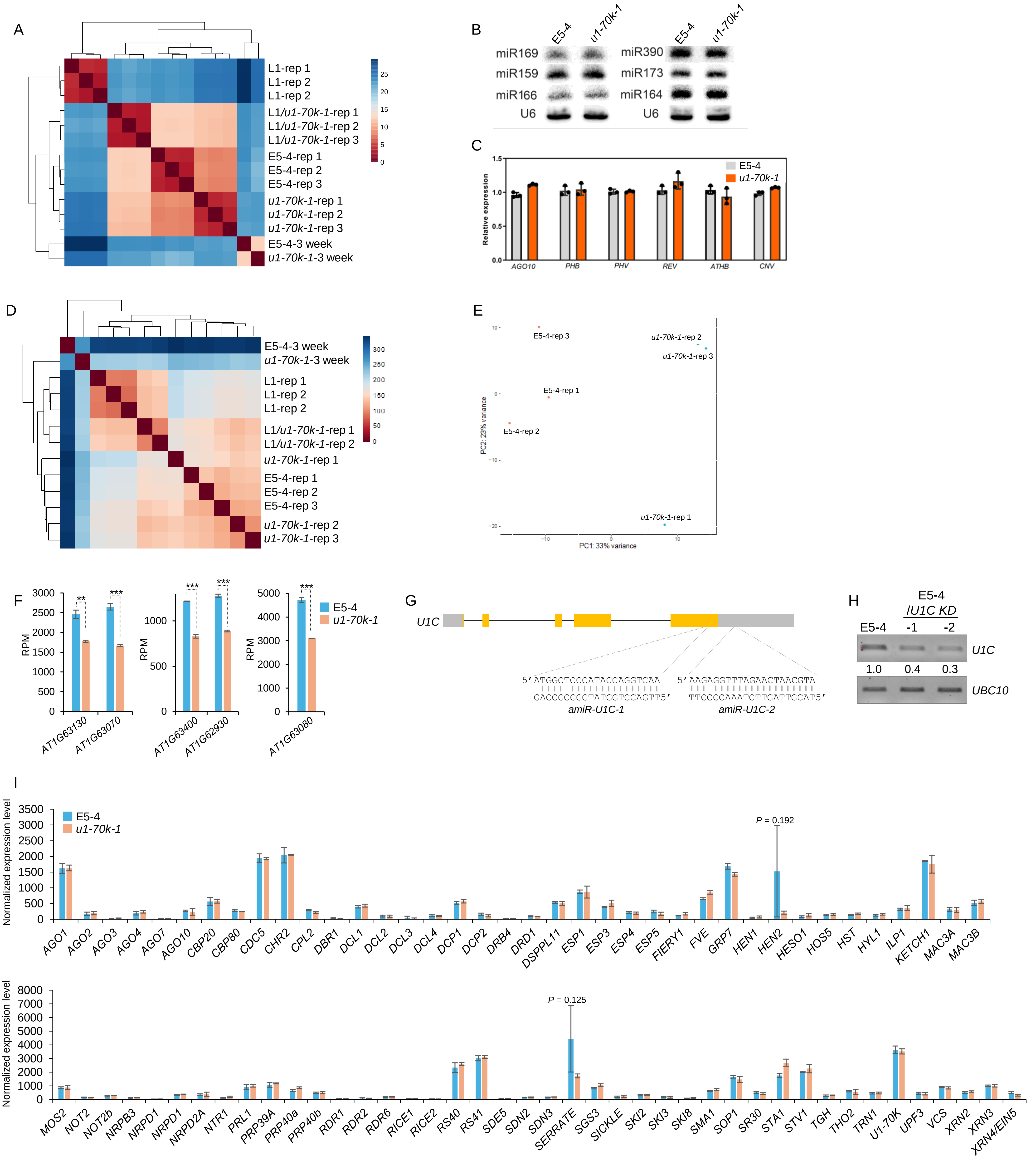
A
L1-rep 1
L1-rep 2
L1-rep 2
L1/u1-70k-1-rep 1
L1/u1-70k-1-rep 2
L1/u1-70k-1-rep 3
E5-4-rep 1
E5-4-rep 2
E5-4-rep 3
u1-70k-1-rep 1
u1-70k-1-rep 2
u1-70k-1-rep 3
E5-4-3 week
u1-70k-1-3 week
B
E5-4
u1-70k-1
miR169
miR159
miR166
U6
miR390
miR173
miR164
U6
C
Relative expression
E5-4
u1-70k-1
AGO10
PHB
PHV
REV
ATHB
CNV
D
E5-4-3 week
u1-70k-1-3 week
L1-rep 1
L1-rep 2
L1-rep 2
L1/u1-70k-1-rep 1
L1/u1-70k-1-rep 2
u1-70k-1-rep 1
E5-4-rep 1
E5-4-rep 2
E5-4-rep 3
u1-70k-1-rep 2
u1-70k-1-rep 3
E
E5-4-rep 3
u1-70k-1-rep 2
u1-70k-1-rep 3
E5-4-rep 1
E5-4-rep 2
u1-70k-1-rep 1
PC2: 23% variance
PC1: 33% variance
F
RPM
E5-4
u1-70k-1
AT1G63130
AT1G63070
AT1G63400
AT1G62930
AT1G63080
G
U1C
5′ ATGGCTCCCATACCAGGTCAA
GACCGCGGGTATGGTCCAGTT5′
amiR-U1C-1
5′ AAGAGGTTTAGAACTAACGTA
TTCCCCAAATCTTGATTGCAT5′
amiR-U1C-2
H
E5-4
/U1C KD
E5-4
-1
-2
U1C
1.0
0.4
0.3
UBC10
I
Normalized expression level
E5-4
u1-70k-1
P = 0.192
AGO1
AGO2
AGO3
AGO4
AGO7
AGO10
CBP20
CBP80
CDC5
CHR2
CPL2
DBR1
DCL1
DCL2
DCL3
DCL4
DCP1
DCP2
DRB4
DRD1
DSPPL11
ESP1
ESP3
ESP4
ESP5
FIERY1
FVE
GRP7
HEN1
HEN2
HESO1
HOS5
HST
HYL1
ILP1
KETCH1
MAC3A
MAC3B
P = 0.125
MOS2
NOT2
NOT2b
NRPB3
NRPD1
NRPD1
NRPD2A
NTR1
PRL1
PRP39A
PRP40a
PRP40b
RDR1
RDR2
RDR6
RICE1
RICE2
RS40
RS41
SDE5
SDN2
SDN3
SERRATE
SGS3
SICKLE
SKI2
SKI3
SKI8
SMA1
SOP1
SR30
STA1
STV1
TGH
THO2
TRN1
U1-70K
UPF3
VCS
XRN2
XRN3
XRN4/EIN5

Fig. S3

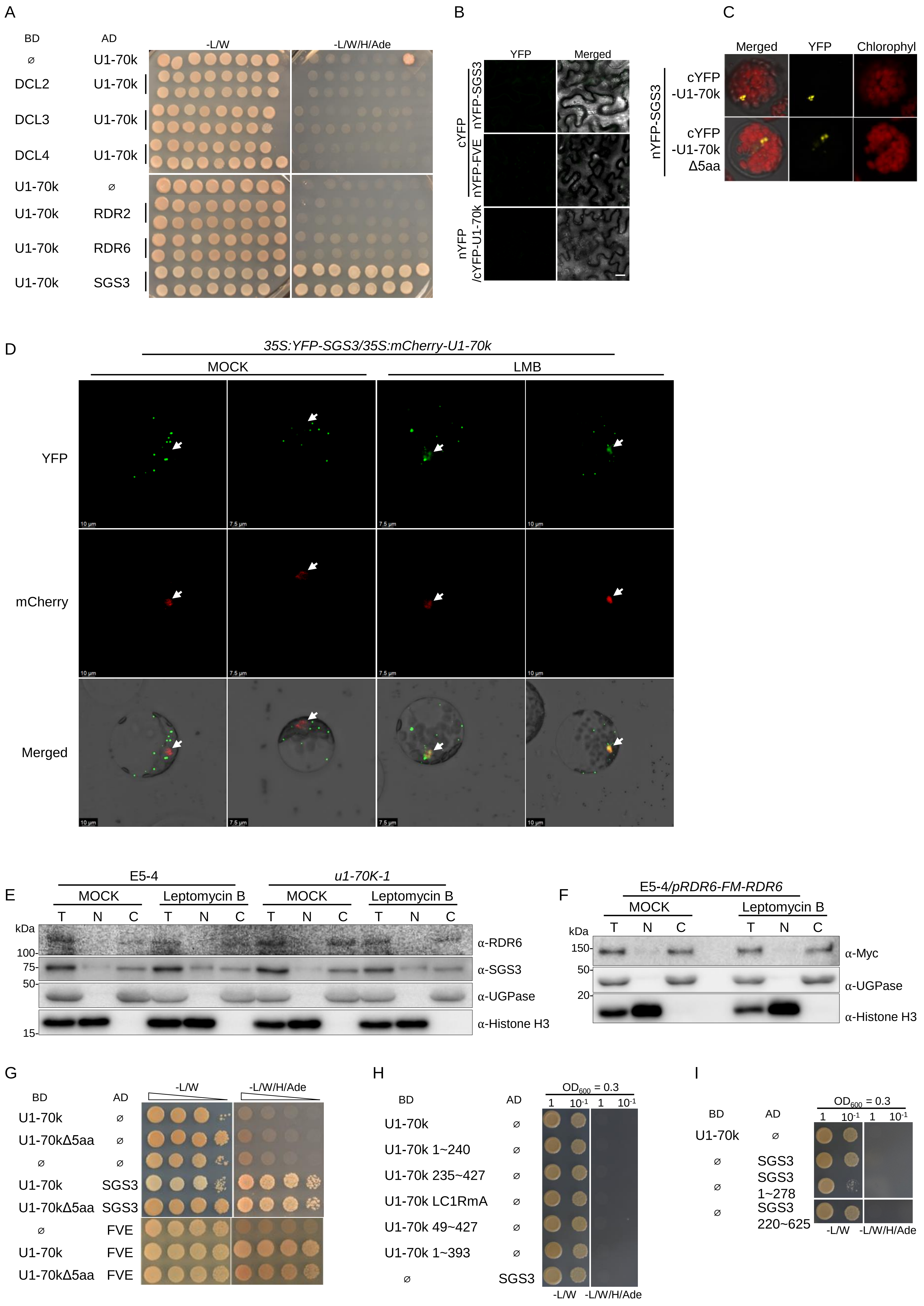

A
BD
AD
-L/W
-L/W/H/Ade
U1-70k
DCL2
DCL3
DCL4
RDR2
RDR6
SGS3
B
YFP
Merged
cYFP
nYFP-SGS3
nYFP-FVE
nYFP
/cYFP-U1-70k
C
Merged
YFP
Chlorophyl
nYFP-SGS3
cYFP
-U1-70k
Δ5aa
D
35S:YFP-SGS3/35S:mCherry-U1-70k
MOCK
LMB
YFP
mCherry
Merged
E
E5-4
u1-70K-1
MOCK
Leptomycin B
T
N
C
kDa
100-
75-
50-
15-
α-RDR6
α-SGS3
α-UGPase
α-Histone H3
F
E5-4/pRDR6-FM-RDR6
150-
20-
α-Myc
G
U1-70kΔ5aa
FVE
H
OD600 = 0.3
1
10-1
U1-70k 1~240
U1-70k 235~427
U1-70k LC1RmA
U1-70k 49~427
U1-70k 1~393
I
SGS3 1~278
SGS3 220~625

### Fig. S4

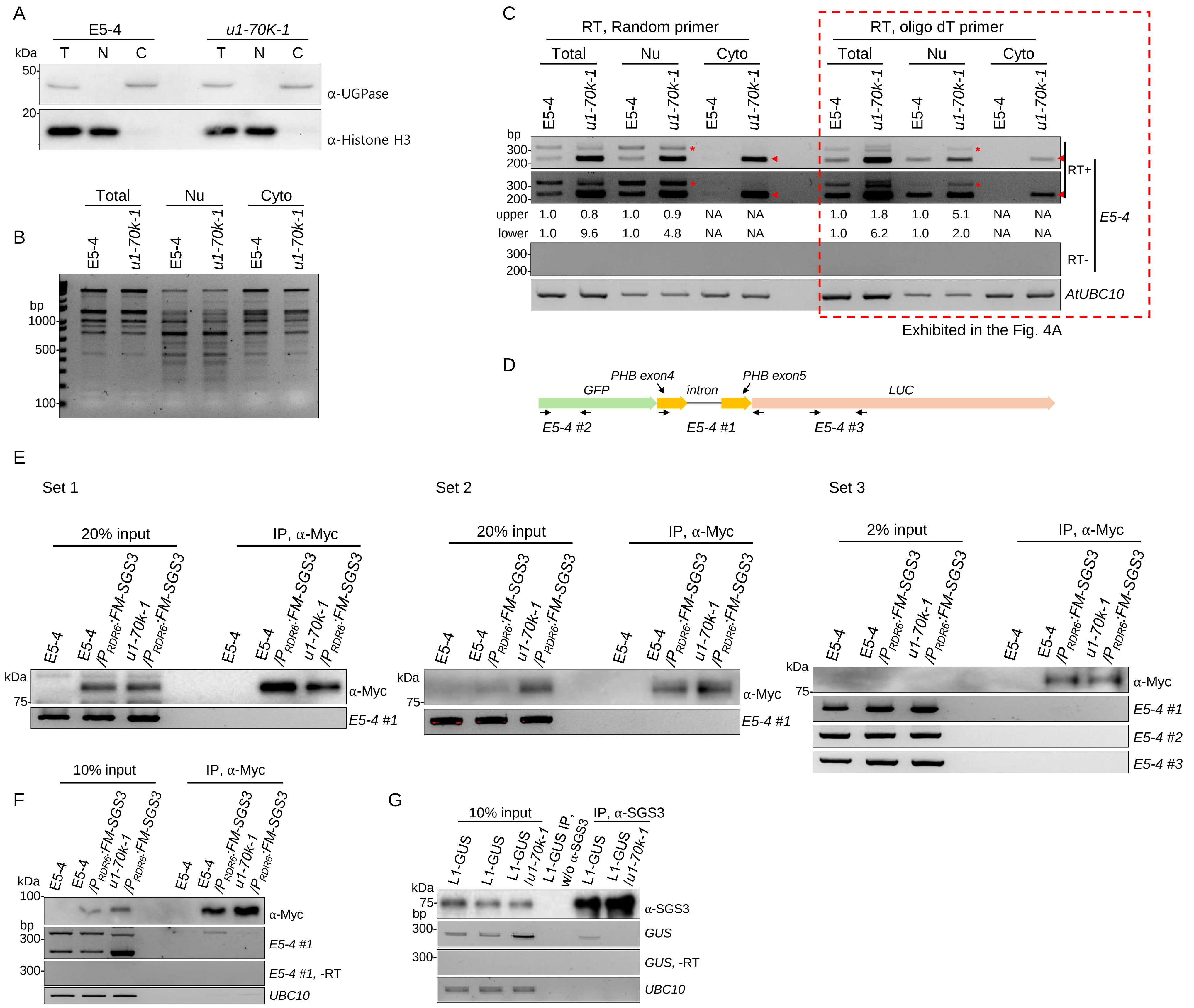
A
E5-4
u1-70K-1
kDa T N C T N C
50
20
α-UGPase
α-Histone H3
B
Total
Nu
Cyto
E5-4
u1-70k-1
bp
1000
500
100
C
RT, Random primer
RT, oligo dT primer
Total
Nu
Cyto
bp
300
200
upper 1.0 0.8 1.0 0.9 NA NA 1.0 1.8 1.0 5.1 NA NA
lower 1.0 9.6 1.0 4.8 NA NA 1.0 6.2 1.0 2.0 NA NA
RT+
RT-
E5-4
AtUBC10
Exhibited in the Fig. 4A
D
PHB exon4
PHB exon5
GFP
intron
LUC
E5-4 #2
E5-4 #1
E5-4 #3
E
Set 1
Set 2
Set 3
20% input
2% input
IP, α-Myc
E5-4
E5-4 /PRDR6:FM-SGS3
u1-70k-1 /PRDR6:FM-SGS3
kDa
75
α-Myc
E5-4 #1
E5-4 #2
E5-4 #3
F
10% input
IP, α-Myc
100
bp
300
E5-4 #1
E5-4 #1, -RT
UBC10
G
10% input
IP, α-SGS3
L1-GUS
L1-GUS /u1-70k-1
L1-GUS IP, w/o α-SGS3
α-SGS3
GUS
GUS, -RT
UBC10

Fig. S5

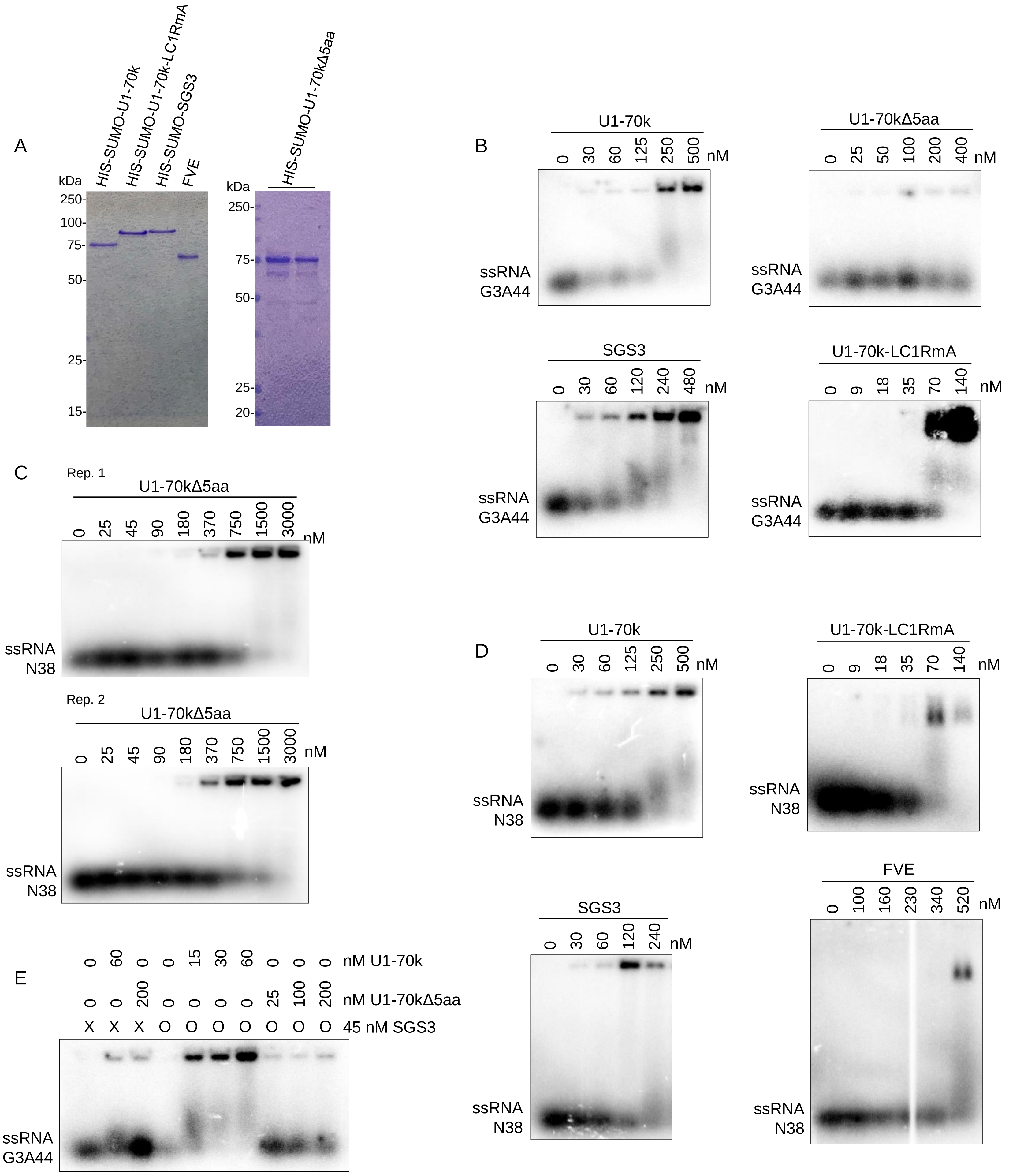

A
kDa
HIS-SUMO-U1-70k
HIS-SUMO-U1-70k-LC1RmA
HIS-SUMO-SGS3
FVE
250-
100-
75-
50-
25-
15-
HIS-SUMO-U1-70kΔ5aa
kDa
250-
75-
50-
25-
20-
B
U1-70k
0 30 60 125 250 500 nM
ssRNA G3A44
U1-70kΔ5aa
0 25 50 100 200 400 nM
ssRNA G3A44
SGS3
0 30 60 120 240 480 nM
ssRNA G3A44
U1-70k-LC1RmA
0 9 18 35 70 140 nM
ssRNA G3A44
C
Rep. 1
U1-70kΔ5aa
0 25 45 90 180 370 750 1500 3000 nM
ssRNA N38
Rep. 2
U1-70kΔ5aa
0 25 45 90 180 370 750 1500 3000 nM
ssRNA N38
D
U1-70k
0 30 60 125 250 500 nM
ssRNA N38
U1-70k-LC1RmA
0 9 18 35 70 140 nM
ssRNA N38
SGS3
0 30 60 120 240 nM
ssRNA N38
FVE
0 100 160 230 340 520 nM
ssRNA N38
E
0 60 0 0 15 30 60 0 0 0 nM U1-70k
0 0 200 0 0 0 0 25 100 200 nM U1-70kΔ5aa
X X X O O O O O O O 45 nM SGS3
ssRNA G3A44

Fig. S6

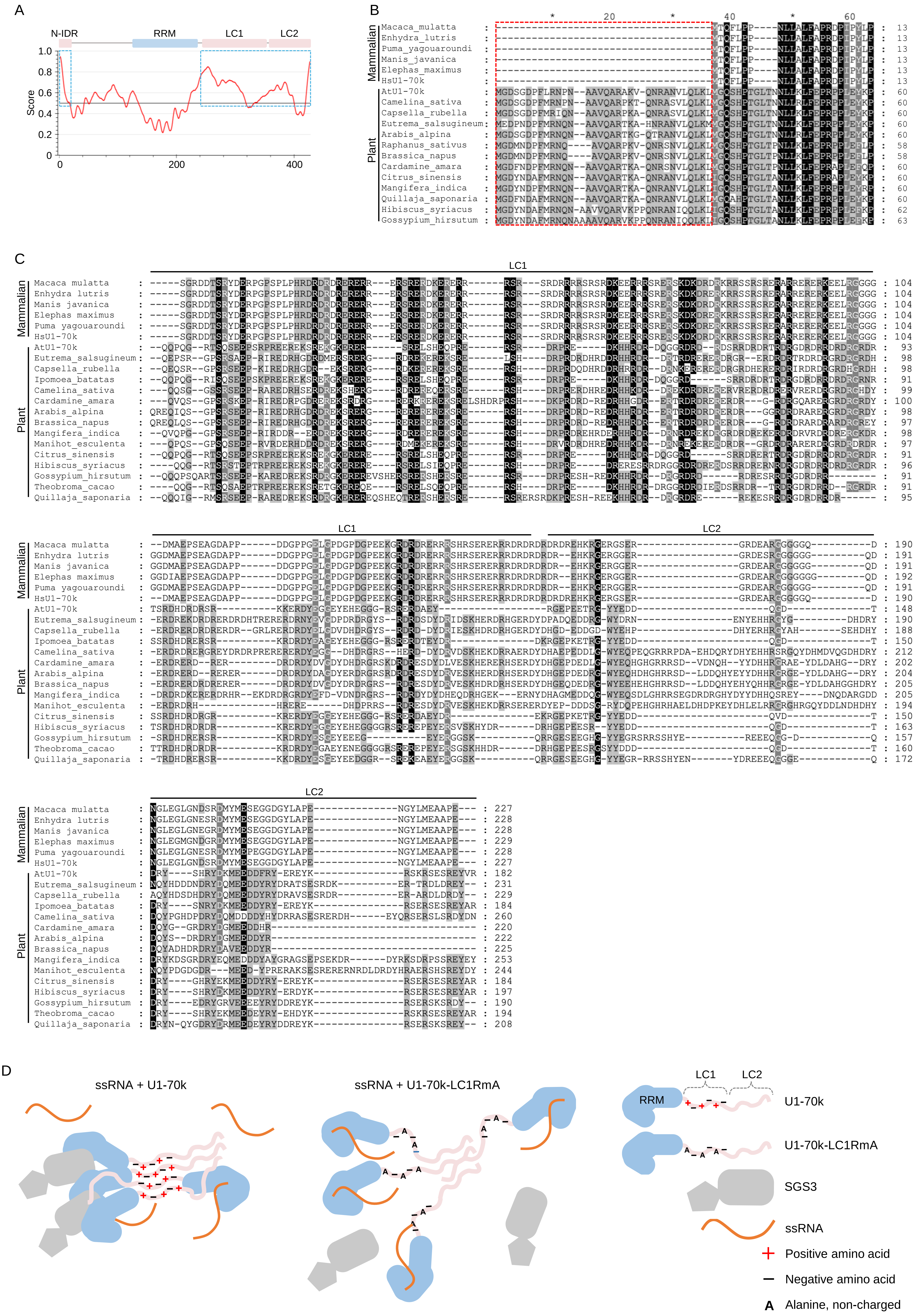
A
N-IDR
RRM
LC1
LC2
Score
1.0
0.8
0.6
0.4
0.2
0
0
200
400
B
Mammalian
Plant
Macaca_mulatta
Enhydra_lutris
Puma_yagouaroundi
Manis_javanica
Elephas_maximus
HsU1-70k
AtU1-70k
Camelina_sativa
Capsella_rubella
Eutrema_salsugineum
Arabis_alpina
Raphanus_sativus
Brassica_napus
Cardamine_amara
Citrus_sinensis
Mangifera_indica
Quillaja_saponaria
Hibiscus_syriacus
Gossypium_hirsutum
C
LC1
LC2
Mammalian
Plant
Macaca mulatta
Enhydra lutris
Manis javanica
Elephas maximus
Puma yagouaroundi
HsU1-70k
AtU1-70k
Eutrema_salsugineum
Capsella_rubella
Ipomoea_batatas
Camelina_sativa
Cardamine_amara
Arabis_alpina
Brassica_napus
Mangifera_indica
Manihot_esculenta
Citrus_sinensis
Hibiscus_syriacus
Gossypium_hirsutum
Theobroma_cacao
Quillaja_saponaria
D
ssRNA + U1-70k
ssRNA + U1-70k-LC1RmA
LC1
LC2
RRM
U1-70k
U1-70k-LC1RmA
SGS3
ssRNA
Positive amino acid
Negative amino acid
A
Alanine, non-charged

Fig. S7

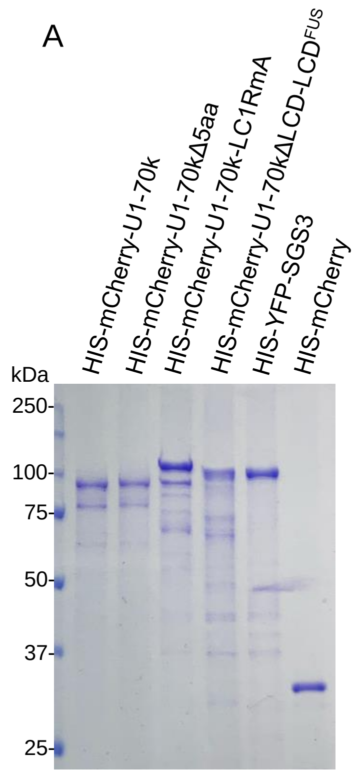

A
HIS-mCherry-U1-70k
HIS-mCherry-U1-70kΔ5aa
HIS-mCherry-U1-70k-LC1RmA
HIS-mCherry-U1-70kΔLCD-LCDFUS
HIS-YFP-SGS3
HIS-mCherry
kDa
250
100
75
50
37
25


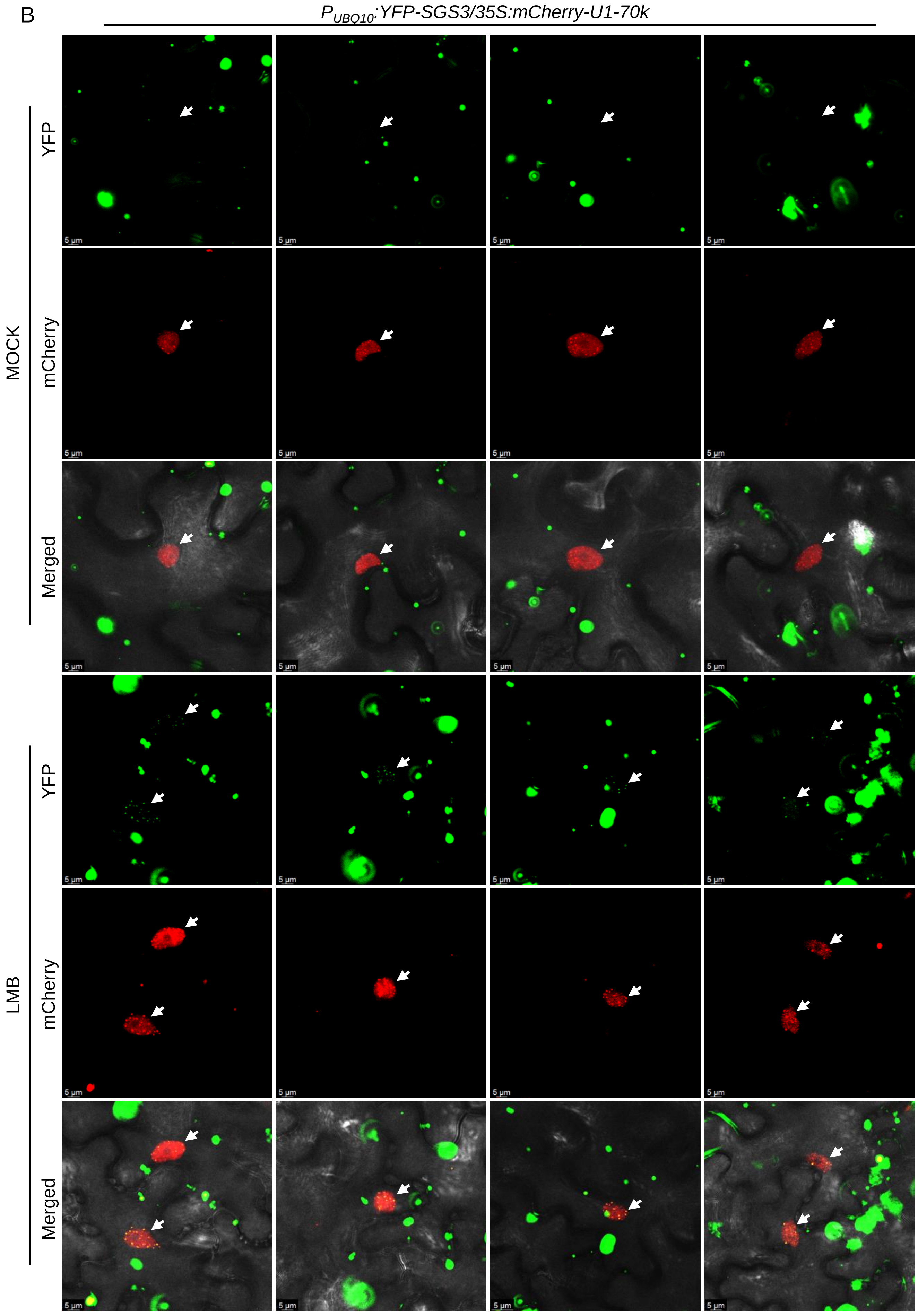

B
PUBQ10:YFP-SGS3/35S:mCherry-U1-70k
MOCK
LMB
YFP
mCherry
Merged
